# Ena/VASP clustering at microspike tips involves Lamellipodin but not I-BAR proteins, and absolutely requires unconventional Myosin-X

**DOI:** 10.1101/2022.05.12.491613

**Authors:** Thomas Pokrant, Jens Ingo Hein, Sarah Körber, Andrea Disanza, Andreas Pich, Giorgio Scita, Klemens Rottner, Jan Faix

**Affiliations:** Institute for Biophysical Chemistry, Hannover Medical School, Carl-Neuberg-Strasse 1, 30625 Hannover, Germany; IFOM ETS - The AIRC Institute of Molecular Oncology, and Department of Oncology and Haemato-Oncology, University of Milan, Milan, Italy; Research Core Unit Proteomics, Hannover Medical School, Carl-Neuberg-Strasse 1, 30625 Hannover, Germany; Division of Molecular Cell Biology, Zoological Institute, Technische Universität Braunschweig, Spielmannstrasse 7, 38106 Braunschweig, Germany; Molecular Cell Biology Group, Helmholtz Centre for Infection Research, Inhoffenstrasse 7, 38124 Braunschweig, Germany

## Abstract

Sheet-like membrane protrusions at the leading edge, termed lamellipodia, drive 2D-cell migration using active actin polymerization. Microspikes comprise actin-filament bundles embedded within lamellipodia, but the molecular mechanisms driving their formation and their potential functional relevance have remained elusive. Microspike formation requires the specific activity of clustered Ena/VASP proteins at their tips to enable processive actin assembly in the presence of capping protein, but the factors and mechanisms mediating Ena/VASP clustering are poorly understood. Systematic analyses of B16-F1 melanoma mutants lacking potential candidate proteins revealed that neither inverse BAR-domain proteins, nor lamellipodin or Abi are essential for clustering, although they differentially contribute to lamellipodial VASP accumulation. In contrast, unconventional myosin-X (MyoX) identified here as proximal to VASP was obligatory for Ena/VASP clustering and microspike formation. Interestingly, and despite the invariable distribution of other relevant marker proteins, the width of lamellipodia in MyoX-KO mutants was significantly reduced as compared to B16-F1 control, suggesting that microspikes contribute to lamellipodium stability. Consistently, MyoX removal caused marked defects in protrusion and random 2D-cell migration. Strikingly, Ena/VASP-deficiency also uncoupled MyoX cluster dynamics from actin assembly in lamellipodia, establishing their tight functional association in microspike formation.

**Significance Statement:** Unlike filopodia that protrude well beyond the cell periphery and are implicated in sensing, morphogenesis and cell-to-cell communication, the function of microspikes consisting of actin-filament bundles fully embedded within lamellipodia is less clear. Microspike formation involves specific clustering of Ena/VASP family members at filament-barbed ends to enable processive actin polymerization in the presence of capping protein, but the factors and mechanisms mediating Ena/VASP clustering have remained unknown. Here, we systematically analyzed these processes in genetic knockout mutants derived from B16-F1 cells and show that Ena/VASP clustering at microspike tips involves Lamellipodin, but not inverse BAR-domain proteins, and strictly requires unconventional Myosin-X. Complete loss of microspikes was confirmed with CRISPR/Cas9-mediated MyoX knockout in Rat2 fibroblasts, excluding cell type-specific effects.

## Introduction

Thin, sheet-like membrane protrusions at the leading edge of cells, known as lamellipodia, drive cell migration in physiological and pathological conditions. The protrusion of the cell front is driven by actin polymerization with the expansion of the polymer directly pushing the membrane forward (1). Actin assembly in the leading edge is driven by the actin-related protein 2/3 (ARP2/3) complex-mediated formation of dendritic actin filament networks downstream of WAVE regulatory complex (WRC) activation and Rac subfamily GTPase signaling (2, 3). The barbed ends of newly formed filament branches are eventually capped by heterodimeric capping protein (CP) (4). However, actin-filament elongation factors such as Ena/VASP or formin family proteins can protect filament ends from capping and also significantly accelerate filament elongation rate (5).

Aside from the branched network, the lamellipodium harbours actin-filament bundles, called microspikes, that span the lamellipodium without protruding beyond the cell edge (6, 7). These structures can display various tilt angles relative to the moving network, the extent of which determines both rates of bundle polymerization and lateral motion (6, 7). Microspikes can convert into filopodia by protruding beyond the cell periphery, and thus separating their polymerization rates from the lamellipodium network (6, 8). However, despite being comprised of parallel actin-filament bundles sharing common constituents, microspikes and filopodia differ considerably. Unlike microspikes, filopodia can form independently of lamellipodial actin networks (9–11). Moreover, filopodia can be formed around the entire cell periphery or the dorsal surface (12–14), and kink, bend or even contribute to the formation of contractile arrays (6). In line with this, ultrastructural analyses recently revealed microspike filaments to be less densely bundled and straight and more bent as compared to filopodial filaments (15).

Ena/VASP proteins are powerful actin polymerases that drive the processive elongation of filament-barbed ends (5, 16). Vertebrates express three paralogs: vasodilator-stimulated phosphoprotein (VASP), mammalian Ena (Mena), and Ena-VASP-like (Evl). All family members harbor an N-terminal Ena/VASP-homology 1 (EVH1) domain essential, but not sufficient, for subcellular localization, followed by a proline-rich region capable of recruiting profilin-actin complexes for actin assembly or interaction with Src-homology 3 (SH3) domain-containing proteins, and a C-terminal EVH2 domain encompassing G-actin and F-actin binding sites and a short coiled-coil motif (Tet) mediating tetramerization (17).

Compared to formins, Ena/VASP tetramers are weakly processive and poorly protect barbed ends from CP in solution (5, 16). However, when clustered at high density on beads, their mode of action is markedly changed allowing them to drive processive and long-lasting actin-filament elongation despite high CP concentrations (5, 18). Ena/VASP proteins were shown to localize to various sites of active actin assembly including the periphery of the leading edge (lamellipodium tip) and the distal tips of actin-bundles termed microspikes and filopodia (17). Consistently, Ena/VASP proteins were previously shown to be important for filopodium formation in *Dictyostelium* and mammalian cells (19, 20). Moreover, contrary to previous assumptions (21), but in agreement with their ability to enhance intracellular motility of *Listeria* (22) or motility of beads in reconstituted systems (23), Ena/VASP proteins have recently been shown to positively regulate cell migration in B16-F1 melanoma cells and fibroblasts (24). Of note, Ena/VASP-deficient B16-F1 cells displayed perturbed lamellipodium architecture and were virtually devoid of microspikes, albeit still capable of forming filopodia (24).

Since Ena/VASP clustering is presumably key for the formation of microspikes, it is instrumental to identify participating factors and determine their specific contributions. According to our current state of knowledge, both insulin receptor substrate of 53 kDa (IRSp53) and lamellipodin (Lpd, also known as Raph1) have been implicated in Ena/VASP clustering (16, 25–27). IRSp53 is a member of the inverted Bin-Amphiphysin-Rvs167 (I-BAR) protein family that senses and/or creates negative membrane curvature by its N-terminal I-BAR domain (also known as IMD), which forms crescent-shaped homodimers (28) that can interact with acidic phospholipids such as PI(4,5)P2 (29). The C-terminal SH3 domain of IRSp53 mediates interactions with proline-rich regions of other proteins such as Ena/VASP members (30) or WAVE2 (31). The remaining four I-BAR protein family members include IRTKS (Baiap2l1), ABBA (MTSS1L), MIM (MTSS1) and Pinkbar (Baiap2l2), albeit the latter appears to be specifically expressed in epithelial cells (32). Notably, IRSp53 was shown to promote recruitment of VASP into leading edge foci in fibroblasts and mediate clustering on beads *in vitro* to initiate VASP-mediated actin assembly in the presence of CP (25). This is supported by very recent work using elaborate *in vitro* reconstitution systems, showing that IRSp53 can recruit and cluster VASP to assemble actin filaments locally on PI(4,5)P2-containing membranes, leading to the generation of actin-filled membrane protrusions resembling filopodia (27).

Lpd on the other hand is a vertebrate member of the Mig-10/RIAM/Lpd (MRL) family of adaptor proteins that localizes to the rim of the leading edge and the tips of microspikes and filopodia (2). The Lpd homodimer is targeted through its Ras-Association and Pleckstrin Homology (RA-PH) domain to the membrane (33), but actin filament binding through highly basic C-terminal sequences lacking the RA-PH domains was also implicated in Lpd recruitment to the leading edge (34). Lpd was assumed to be obligatory for Ena/VASP protein recruitment by its 6 EVH1-binding motifs to the leading edge to promote lamellipodia protrusion (33, 34), but its genetic elimination challenged this view (35). The actin filament-binding activity of Lpd, which can occur independently of Ena/VASP binding, was proposed to trigger Ena/VASP clustering and tether the complexes to growing barbed ends, thereby increasing their processive polymerase activity (34). More recent work proposed the initial formation of small leading edge-Lpd clusters devoid of IRSp53 that expand, fuse and subsequently recruit VASP to induce filopodia/microspike formation in B16-F1 cells (26).

To resolve the controversy on which proteins are required for VASP clustering at the leading edge and for microspike formation, we employed CRISPR/Cas9-technology combined with proteomics to systematically analyze VASP clustering and microspike formation in genetic mutants derived from B16-F1 cells, which have emerged as excellent model system for the dissection of lamellipodium protrusion and microspike formation.

## Results

### Generation of a B16-F1-derived mutant cell line devoid of all I-BAR proteins

To investigate the function of the I-BAR family in cell migration, we first employed semi-quantitative, reverse transcription polymerase chain reaction (RT-PCR) to compare expression levels of the 5 known family members in B16-F1 mouse melanoma cells (SI Appendix, Figs. S1*A* and *B*). Compared to control PCRs, conducted with plasmid DNA templates harboring respective full-length cDNAs, we could amplify IRSp53 and IRTKS at high and comparable levels from a B16-F1 cDNA library. ABBA and MIM, on the other hand, were amplified only at low or very low levels, respectively, while the epithelial-specific Pinkbar was not expressed. Consistent with previous observations (36–39), ectopic expression of IRSp53, IRTKS, ABBA and MIM fused to EGFP showed accumulation of the fusion proteins at the tips of protruding lamellipodia and at intracellular vesicles (SI Appendix, Fig. S1*C*). These findings prompted us to employ CRISPR/Cas9 technology and sequentially inactivate the four respective genes to obtain a B16-F1 mutant cell line devoid of all expressed I-BAR family members. Successful, consecutive inactivation of IRSp53 (designated as 1xKO) and IRSp53/IRTKS (2xKO) in independent clonal cell lines was confirmed by TIDE sequence trace decomposition analyses (40), and further validated by immunoblotting (SI Appendix, Fig. S1*D*). Subsequent elimination of ABBA in the 2xKO mutant yielded the IRSp53/IRTKS/ABBA 3xKO mutant, and the 4xKO mutant devoid of IRSp53/IRTKS/ABBA/MIM was obtained by elimination of MIM in the 3xKO mutant. Successful disruption of the *Mtss2* and *Mtss1* genes encoding ABBA and MIM, respectively, was confirmed by TIDE analysis and sequencing of genomic target sites (SI Appendix, Fig. S1E).

### Loss of I-BAR proteins impairs 2D random cell migration and lamellipodia formation

Migration rates of B16-F1 wild-type and mutant cells of each genotype were then analyzed on laminin by phase-contrast, time-lapse microscopy. Interestingly, consecutive removal of the most highly expressed IRSp53 and IRTKS caused an increasing phenotype in 2D-migration rate, ranging from a modest reduction to 1.25±0.36 μm min^-1^ in single IRSp53-KO mutants (1xKO) as compared to control (1.44±0.42 μm min^-1^; mean±SD) to 1.00±0.33 μm min^-1^ in the IRSp53/IRTKS double-KO (2xKO) (Fig. 1*A*). Additional elimination of ABBA and MIM, on the other hand, had virtually no further effect on cell speed as compared to the double mutant, with 1.03±0.31 μm min^-1^ in the 3xKO and 1.04±0.34 μm min^-1^ in the 4xKO (Fig. 1 *A*). This was accompanied by a gradual, but moderate increase in directionality, again culminating in 2xKO mutant cells that were about 20% more directional as compared to the wild-type (SI Appendix, Fig. S2). Finally, we calculated the mean square displacement (MSD) in wild-type cells and the entire collection of mutants to assess their effective directional movement. In spite of their slightly higher directionality, presumably owing to their lower speed, the 1xKO and 2xKO I-BAR-mutant cells displayed incrementally decreasing and lower MSD values as compared to wild-type, while 3xKO and 4xKO were again indistinguishable from the 2xKO mutant (Fig. 1*B*).

**Fig. 1.**
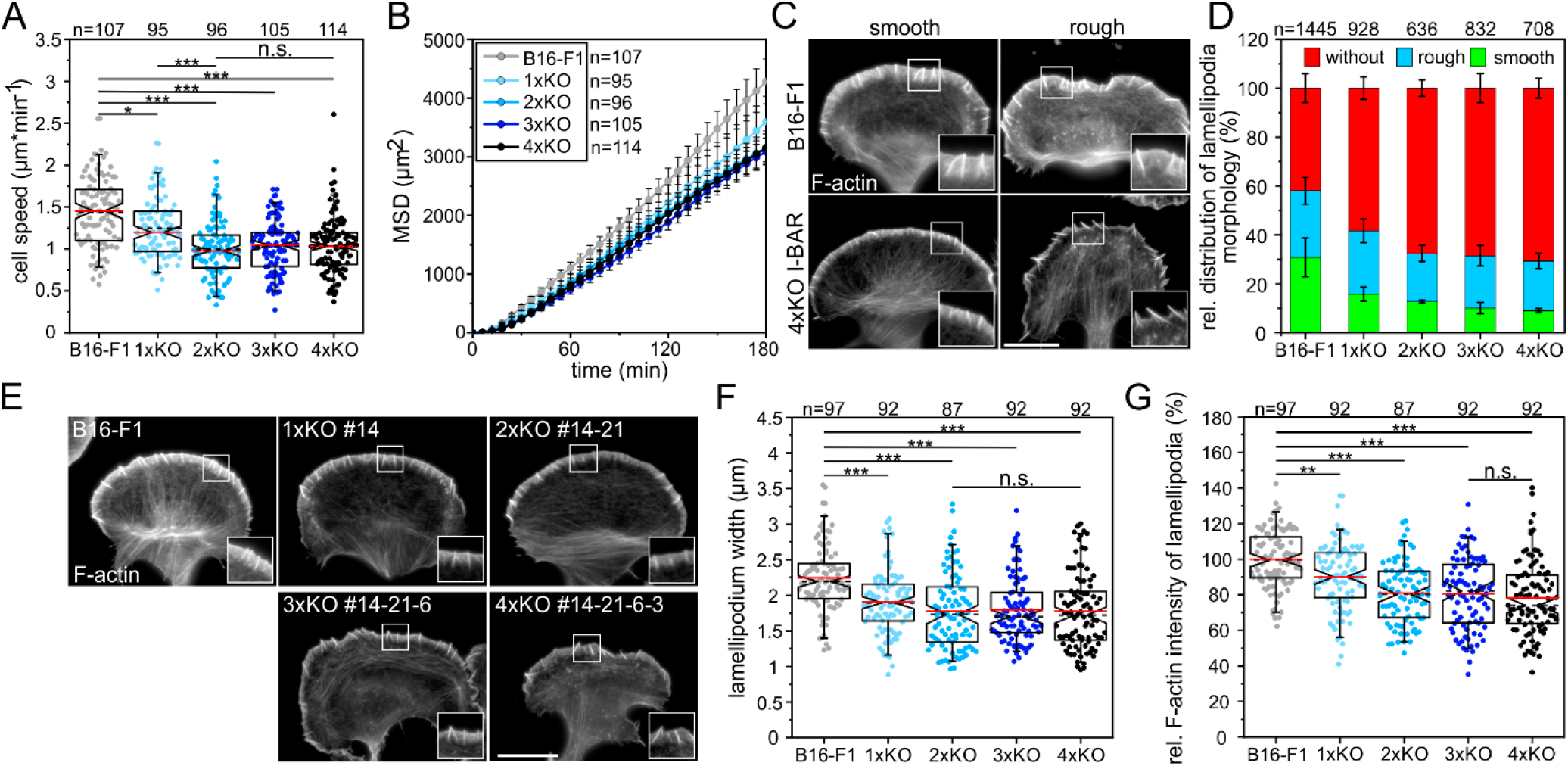
Loss of I-BAR proteins impairs 2D-cell migration and affects lamellipodial dynamics in B16-F1 cells. (*A*) Consecutive gene disruption of the two most highly expressed I-BAR family members IRSp53 and IRTKs increasingly diminished cell migration on laminin, whereas additional loss of ABBA and MIM were negligible. (*B*) Analyses of mean square displacement of wild-type versus mutant cells. Error bars represent means±SEM. (*C*) Morphologies of lamellipodia in B16-F1 and 4x-KO I-BAR mutant cells. Cells migrating on laminin were stained for F-actin with phalloidin. Bar, 15 μm. (*D*) Quantification of lamellipodia formation in wild-type and mutant cells. Bars represent arithmetic means±SD. (*E*) Representative examples of lamellipodia in B16-F1 cells and derived I-BAR mutants after phalloidin staining. Bar, 15 μm. (*F* and *G*) Quantification of lamellipodium widths and F-actin intensities in lamellipodia of wild-type and mutant cells. (C and E) Insets, enlarged images of boxed regions. (A, F, G) Notched boxes in box plots indicate 50% (25-75%) and whiskers 90% (5-95%) of all measurements. Arithmetic means are highlighted in red and dashed black lines depict medians. (A, F) Kruskal-Wallis test or (G) 1-way ANOVA were used for statistical analysis. *p≤0.05, **p≤0.01, ***p≤0.001; n.s.; not significant. n, number of cells. A-B, D, F-G show pooled data from at least three independent experiments.

Most reports on B16-F1 cells focus on a subpopulation of migrating cells harboring prominent lamellipodia. Although time-lapse video microscopy data prove that virtually every B16-F1 wild-type cell is capable of forming productive lamellipodia, the percentage of cells with fully-developed lamellipodia will only correspond to app. 40% at any given time (41). To test how I-BAR proteins contribute to lamellipodia shapes, we compared morphologies of wild-type and all I-BAR-mutant clones focusing on their leading edge morphologies, which we here classified as smooth or rough by imaging after phalloidin staining (Fig. 1*C*). In contrast to wild-type, in which 30.8±7.9% of the cells formed smooth lamellipodia according to this categorization, this proportion was gradually decreased by 70% to 9.1±0.7% in 4xKO cells (Fig. 1*D*). Notably, I-BAR protein removal mostly increased the cell fraction lacking lamellipodia (without), but only slightly affected the percentage of cells with rough edges (Fig. 1*D*), suggesting that I-BAR proteins particularly optimize the formation of smooth lamellipodia.

Since 2D random cell migration on flat surfaces is largely driven by lamellipodial actin assembly in B16-F1 cells, we then analyzed actin-filament (F-actin) content in wild-type and I-BAR KO cells after phalloidin staining. In contrast to controls, which displayed prominent lamellipodia containing numerous microspikes, I-BAR KO cells developed increasingly compromised lamellipodia that somewhat unexpectedly though still formed microspikes (Fig. 1*E* and see below). The average width of lamellipodia in the quadruple mutant was reduced by about 21% to 1.8±0.5 μm when compared to control (2.2±0.5 μm), and lamellipodial actin-filament density was similarly reduced (Figs. 1*F* and *G*). Finally, we asked to which extent lamellipodia protrusion was affected. To this end, we recorded wildtype and I-BAR-deficient cells randomly migrating on laminin by time-lapse, phase-contrast microscopy, and determined respective protrusion rates by kymograph analyses (SI Appendix, Figs. S3*A* and *B*). Quantification revealed lamellipodia protrusion to be less coordinated and reduced by about 20% in the quadruple mutant as compared to control (SI Appendix, Figs. S3*C* and Movie S1). In addition to reduced protrusion, lamellipodia persistence was also diminished by roughly 25% (SI Appendix, Figs. S3*D*), strongly suggesting that I-BAR family proteins capable of contributing to negative membrane curvatures at cell edges significantly contribute to the efficacy and stability of lamellipodial protrusions.

### I-BAR proteins are dispensable for the formation of microspikes and filopodia

IRSp53 and IRTKs have been assumed to be essential regulators of filopodia formation, since they couple Rho-GTPase signaling to actin cytoskeleton and membrane remodeling (30, 42, 43). Hence, we analyzed microspike and filopodia formation in the I-BAR mutants in more detail. Notably, microspikes, which are usually fully embedded into lamellipodia in B16-F1 cells, frequently protruded in an unusual manner beyond the plane of the plasma membrane in the 4xKO mutant (SI Appendix, Fig. S4A). Quantification further revealed microspikes to be statistically significantly longer in 4xKO mutant (3.17±0.94 μm) as compared to B16-F1 control (2.89±0.69 μm) cells (SI Appendix, Fig. 4B). Thus, considering their narrower lamellipodia (Fig. 1*F*), the quadruple mutant formed microspikes that were almost twice as long as the lamellipodia they are associated with, whereas microspikes in wild-type cells were comparable in length to the width of their lamellipodia (SI Appendix, Fig. 4C). Surprisingly, however, the quadruple mutant showed no noticeable change in the average number of microspikes per cell as compared to control (SI Appendix, Fig. 4D). This finding was unanticipated, as each microspike bundle should harbor a VASP cluster at its tip to allow processive actin-filament elongation in the presence of CP. Since prominent VASP clusters were still formed at the membrane even in the absence of I-BAR proteins (SI Appendix, Fig. 4E), these factors alone are evidently not mandatory for the formation of microspikes and the clustering of Ena/VASP proteins at their tips. To our surprise, the majority of 4xKO cells devoid of prominent lamellipodia also formed numerous filopodia as evidenced by localization of VASP at their tips (SI Appendix, Fig. 4E). Thus, to further assess whether loss of I-BAR function affects the formation of filopodia, we first explored B16-F1 and 4xKO cells after ectopic expression of EGFP-tagged, full-length myosin-X (MyoX) or the constitutively active variant of the formin mDia2 (mDia2ΔDAD), both of which are well known to induce filopodia (12, 44, 45) (SI Appendix, Fig. S5A). Notably, quantification of induced filopodia revealed no significant difference between the quadruple mutant as compared to wild-type cells (SI Appendix, Fig. S5B), suggesting that I-BAR proteins are also dispensable for filopodia formation in B16-F1 cells. We then induced filopodia in the absence of lamellipodia in B16-F1 and 4xKO I-BAR cells by inhibiting Arp2/3 complex with high concentrations of CK666 combined with cell seeding on low laminin concentration (24). Analyses of phalloidin-stained cells revealed 4xKO I-BAR mutant cells forming more than twice the number of filopodia compared to control (SI Appendix, Figs. S5C and D). To test whether this effect was cell-type specific, we then consecutively eliminated the two most abundant I-BAR proteins IRSp53 and IRTKs in NIH 3T3 fibroblasts by CRISPR/Cas9 technology (SI Appendix, Figs. S5E and F). Since filopodia formation is rare in fibroblasts when migrating on high fibronectin, we again treated these cells with CK666 on low fibronectin concentrations. In this case, analyses of wild-type and 2xKO mutant fibroblasts showed no marked differences in filopodium formation (SI Appendix, Figs. S5G and H and Movie S2), suggesting that the enhanced filopodia formation after loss of I-BAR proteins is cell-type specific. Nevertheless, these data further indicate that I-BAR proteins are dispensable for filopodia formation.

### Loss of Lpd perturbs microspike formation

An alternative potential candidate implicated in VASP clustering at the lamellipodial edge is Lamellipodin (Lpd), since it was reported to form small leading-edge clusters that subsequently recruit VASP and initiate actin assembly (26). Notably, both VASP and Lpd also bind filamentous actin (34), suggesting that clustering might even be accomplished by shared interactions between these three proteins. However, the genetic disruption of Lpd has not eliminated Ena/VASP targeting to the leading edge of lamellipodia (35). This notwithstanding, the formation of microspikes and VASP clustering in Lpd-deficient cells have hitherto not been analyzed. Thus, to explore potential functions of Lpd and VASP in these processes, we first expressed and purified previously characterized dimeric GCN4-Lpd^850-1250^ (WT) and its actin-binding deficient variant GCN4-Lpd^850-1250^ (44A) (34), additionally carrying an N-terminal SNAP-tag. After coupling of the proteins to SNAP-capture magnetic beads, we monitored actin assembly in the absence or presence of soluble VASP and CP by TIRF imaging (Fig. 2*A*). Unexpectedly, Lpd-WT was already sufficient to initiate weak actin assembly in the absence of VASP on the bead surface (Fig. 2*B*). In contrast, VASP addition resulted in strong actin assembly on the surface of Lpd-derivatized beads, with the filaments elongating about three times faster in the presence of the Lpd-44A mutant as compared to Lpd-WT (Fig. 2*B*, SI Appendix, Fig. S6 and Movie S3). This shows not only that both Lpd-constructs are very effective to recruit VASP from solution and form clusters on the bead surface to initiate processive actin-filament elongation in the presence of CP, but in addition that the F-actin binding site in Lpd is dispensable and counterproductive for actin assembly, at least *in vitro*.

**Fig. 2.**
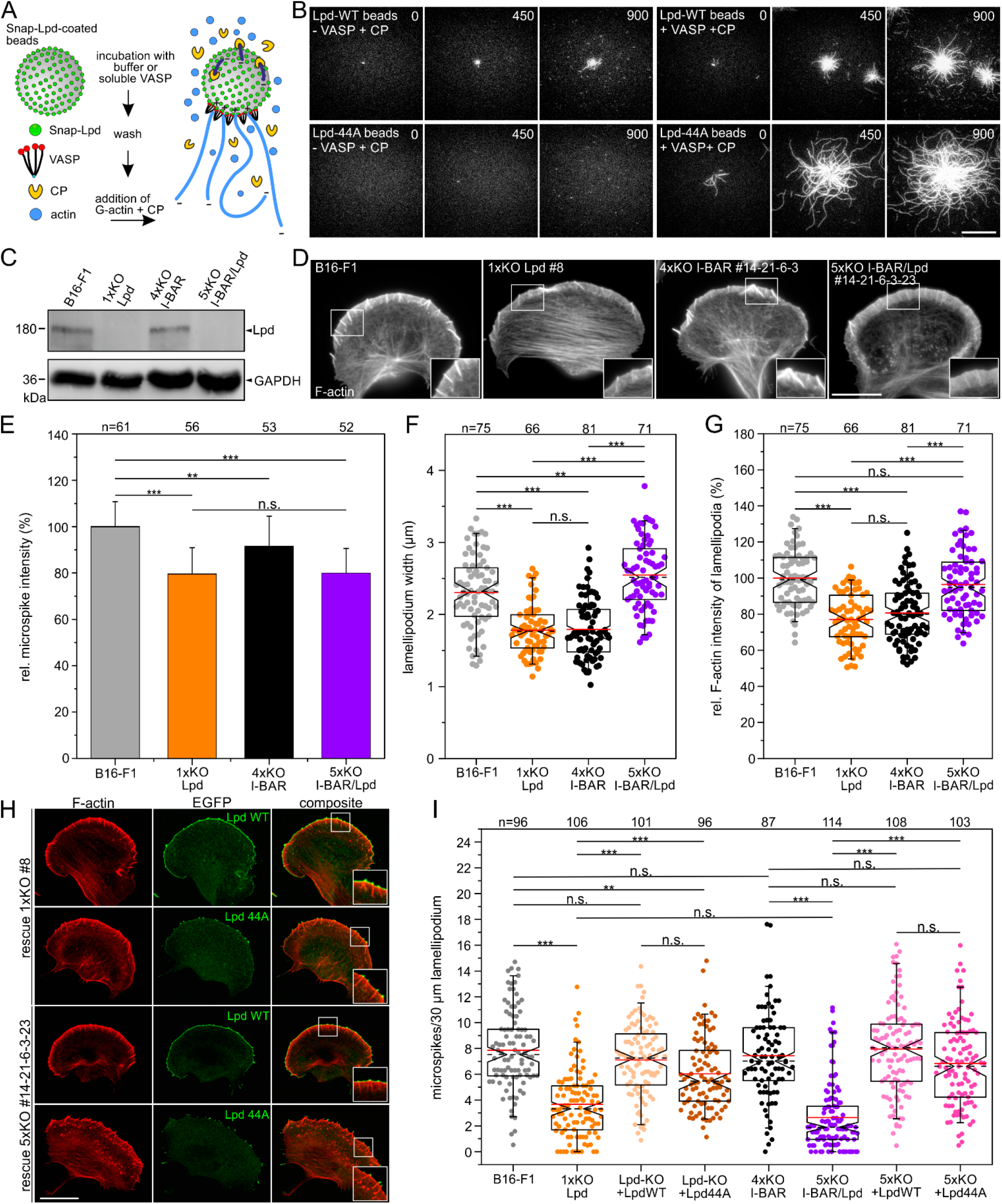
Lpd contributes to microspike formation. (A) Scheme of TIRF-M assays analyzing VASP-mediated actin assembly on SNAP-Lpd-derivatized beads in the presence of capping protein (CP). (*B*) Still images from time-lapse TIRF-movies using 1 μM actin (10% Atto488-labelled) together with 25 nM CP and either wild-type Lpd^850-1250^ (Lpd-WT) or mutant Lpd^850-1250^ (Lpd44A) on derivatized beads pre-incubated with buffer or soluble VASP. Bar, 10 μm. (*C*) Loss of Lpd in 1xKO Lpd or 5xKO I-BAR/Lpd mutant cells was confirmed by immunoblotting. GAPDH was used as loading control. (*D*) Representative images of wild-type and mutant cells as indicated and stained for F-actin. Bar, 15 μm. (*E*) Quantification of microspike intensities in wild-type and mutants cells. Bars represent means±SD. (*F*) Quantification of lamellipodium width. (*G*) Quantification of F-actin intensity. (*H*) Positively charged residues in the C-terminus of Lpd are critical for membrane targeting. Representative images of 1xKO Lpd and 5xKO I-BAR/Lpd mutant cells reconstituted with full-length Lpd-WT *versus* Lpd-44A fused to EGFP. Fixed cells were stained for the F-actin and EGFP signals enhanced with nanobodies. Bar, 2O μm. (*I*) Quantification of microspikes in wild-type, mutant and reconstituted mutant cells. (D and H) Insets, enlarged images of boxed regions. (E-G, I) Notched boxes in box plots indicate 50% (25-75%) and whiskers 90% (5-95%) of all measurements; arithmetic means are highlighted in red and dashed black lines depicting the medians. (E-F, I) Kruskal-Wallis test or (G) 1-way ANOVA were used for statistical analysis. *p≤0.05, **p≤0.01, ***p≤0.001; n.s.; not significant. n, number of cells. E-G and I show pooled data from three independent experiments.

Given the conceivable role of Lpd in microspike formation and VASP clustering, possibly in conjunction with I-BAR-family proteins, we eliminated Lpd by CRISPR/Cas9 in independent quadruple mutants. Protein elimination in a representative set of mutants used for further detailed analyses was confirmed by TIDE analyses and immunoblotting (Fig. 2*C*). We then analyzed F-actin distribution in parental, previously described 1xKO Lpd clone #8 (35) and the newly generated 5xKO I-BAR/Lpd mutant #14-21-6-3-23 after phalloidin staining (Fig. 2*D*). Compared to B16-F1 control cells, which exhibited prominent lamellipodia with numerous microspikes, Lpd-KO cells developed regular lamellipodia largely consistent with previous findings (35), except that they appeared to form fewer microspikes. Unexpectedly, and as opposed to the commonly rough and more chaotic nature of the I-BAR quadruple mutant, combined loss of Lpd- and I-BAR-functions in the 5xKO mutant led to exaggerated and exceptionally smooth lamellipodia, which contained fewer microspikes (Movie S4). Notably, F-actin content in microspikes was reduced by about 20% in Lpd-KO and 5xKO I-BAR/Lpd mutants compared to wild-type, while it was more moderately affected (reduction by 10%) in 4xKO cells, suggesting microspike bundles to contain fewer actin filaments particularly after loss of Lpd (Fig. 2*E*). Furthermore, the average width of lamellipodia was even increased in the 5xKO I-BAR/Lpd mutant by about 10% to 2.5±0.5 μm as compared to B16-F1 control with 2.3±0.5 μm, while it was reduced by 23% (1.8±0.3 μm) in the Lpd-KO or 22% with (1.8±0.4 μm) in 4xKO I-BAR mutant, respectively (Fig. 2*F*). Quantification of F-actin content in lamellipodia revealed no changes in the 5xKO when compared to wild-type, while the levels in 1xKO Lpd and in 4xKO I-BAR mutant cells were found here to be reduced by 23% and 19%, respectively (Fig. 2*G*).

To better understand these complex lamellipodial phenotypes, we then stained the cells for the marker proteins VASP, WAVE2 and Arp2/3 complex. Notably, despite comparable expression levels, VASP accumulation in the tips of lamellipodia of the 5xKO I-BAR/Lpd mutant was again comparable to wild-type, while it was reduced by almost 25% in both the 1xKO Lpd and the 4xKO I-BAR mutants (SI Appendix, Figs. S7A-C). WAVE2, on the other hand, was only diminished by about 25% in lamellipodia in 4xKO I-BAR as compared to the other cell lines (SI Appendix, Figs. S7D-F). Unexpectedly, the expression levels of WAVE2 in the 4xKO I-BAR and 5xKO I-BAR/Lpd mutant were reduced by about 20% as compared to the other cell lines as assessed by immunoblotting and densitometry, explaining diminished accumulation in lamellipodia of the 4xKO I-BAR mutant, but not in the 5xKO I-BAR/Lpd mutant. Interestingly, Arp2/3 complex accumulated in the 5xKO mutant in a peripheral band even broader than observed in wild-type lamellipodia (increased by 14%), but with roughly 17% lower intensity, whereas Lpd-KO and 4xKO cells exhibited 20% narrower bands as compared to wild-type (SI Appendix, Figs. S7G-I). These rather unexpected findings were validated by comparable analyses with a full set of independent mutants to exclude potential off-target effects during somatic genome editing (SI Appendix, Figs. S8). This was corroborated by a side-by-side comparison of the expression levels of relevant lamellipodial marker proteins from all these mutants and revealed that only the WRC components WAVE2 and Abi1 were specifically reduced to a similar extent in the independent clonal cell lines after I-BAR protein removal (SI Appendix, Fig. S9). Thus, despite lower WAVE2 expression, the combined loss of Lpd and I-BAR functions apparently alleviates defects in lamellipodium formation, at least in part, by improved VASP and WAVE recruitment as compared to the isolated I-BAR or Lpd-KO conditions.

Since loss of Lpd exhibited the greatest impact on microspike formation both with and without I-BAR family members, we finally compared the ability of EGFP-tagged, full-length Lpd-WT and Lpd-44A constructs to rescue microspike formation in Lpd-KO and I-BAR/Lpd-KO cells (Fig. 2*H*). Although both constructs could rescue microspike formation reasonably well, only Lpd-WT accumulated prominently both at the tips of microspikes and the leading edge. By contrast, Lpd-44A was only detected in microspike tips, but failed to accumulate along the lamellipodial cell front. In line with diminished recruitment of Lpd-44A to the protruding front, the extent of microspike rescue in reconstituted Lpd-KO and 5xKO I-BAR/Lpd mutant cell lines revealed the rescue to be more effective with the Lpd-WT construct (Fig. 2*I*). Interestingly, the combined elimination of Lpd and the I-BAR family proteins resulted in a stronger defect in microspike formation as compared to the Lpd-KO mutant, suggesting that both protein families synergize to some extent in this process, although Lpd clearly has the greater impact as compared with I-BAR proteins. Considering that microspikes always harbor a VASP cluster at their tips, these rescue experiments of Lpd-KO and 5xKO I-BAR/Lpd cells were also confirmed to restore VASP cluster formation in a comparable manner (SI Appendix, Fig. S10). Taken together, all these results show that Lpd contributes, but is not essential for microspike formation and VASP clustering in the lamellipodium.

### Abi recruits VASP to the leading edge but is dispensable for clustering

Staining 5xKO I-BAR/Lpd mutant cells for endogenous VASP, we detected prominent clusters of the protein in the leading edges of these cells, clearly suggesting the existence of factors additional to the I-BAR family and Lpd apparently operating in Ena/VASP clustering (Fig. 3*A*). Notably, the latter were previously shown to directly interact with the WRC component Abi in different cell types (46–48). Moreover, activated WRC was reported to form large, oligomeric complexes in protruding regions of the plasma membrane (49), suggesting that Abi might contribute to Ena/VASP clustering at microspike tips. Since Abi binds by virtue of proline-rich stretches in its C-terminus to the EVH1-domain of Ena/VASP (48), we generated and purified artificially-dimerized SNAP-tagged fragments of WT (Abi1^331-424^) and an Abi mutant (Abi1^331-424^-mut), in which the critical proline residues were replaced by alanines (Fig. 3*B*). Additionally, we inactivated the Abi1-SH3 domain because Ena/VASP proteins can also interact with SH3-domains (30). After coupling of the proteins to SNAP-capture magnetic beads, we again monitored actin assembly in the absence or presence of soluble VASP and CP by TIRF imaging (Fig. 3*C* and Movie S5). Abi1-WT promoted recruitment of VASP followed by prominent actin assembly on the bead surface in the presence of CP, whereas filament growth was not detectable on Abi1-mut-derivatized beads, suggesting that Abi could be involved in Ena/VASP clustering also in cells. To explore this potential role of Abi, we then simultaneously eliminated Abi1 and Abi2 by CRISPR/Cas9 in the 5xKO I-BAR/Lpd cells yielding 7xKO I-BAR/Lpd/Abi mutants. Loss of both proteins was confirmed by immunoblotting (Fig.3*D*). Consistent with previous work (10, 41, 50), loss of functional WRC in independent 7xKO mutants was associated with complete loss of canonical lamellipodia (SI Appendix, Fig. S11). Reconstitution of 7xKO cells with EGFP-tagged, full-length Abi1-WT and Abi1-mut revealed that both constructs efficiently rescued lamellipodium formation as assessed by staining for the WRC component WAVE2 (Figs. 3*E* and *F*). However, only cells expressing EGFP-Abi1-WT were able to efficiently recruit VASP to the leading edge, whereas VASP accumulation in Abi1-mut expressing cells was diminished by about 84% (Figs. 3*E* and *F*). The dependence of VASP accumulation in pseudopods or mammalian cell lamellipodia on the presence of WRC was already previously established using complementary experimental approaches (47, 51), but the data shown here confirm a major role in this process for the WRC subunit Abi and its proline-rich domains. In contrast, however, in cells expressing EGFP-Abi1-mut, VASP was still capable of associating in prominent clusters at microspike tips (Fig. 4*E* and SI Appendix, Fig. S12), suggesting that while being required for VASP recruitment to the lamellipodium edges in between microspikes, its recruitment to and clustering at microspike tips can occur independently of Abi.

**Fig. 3.**
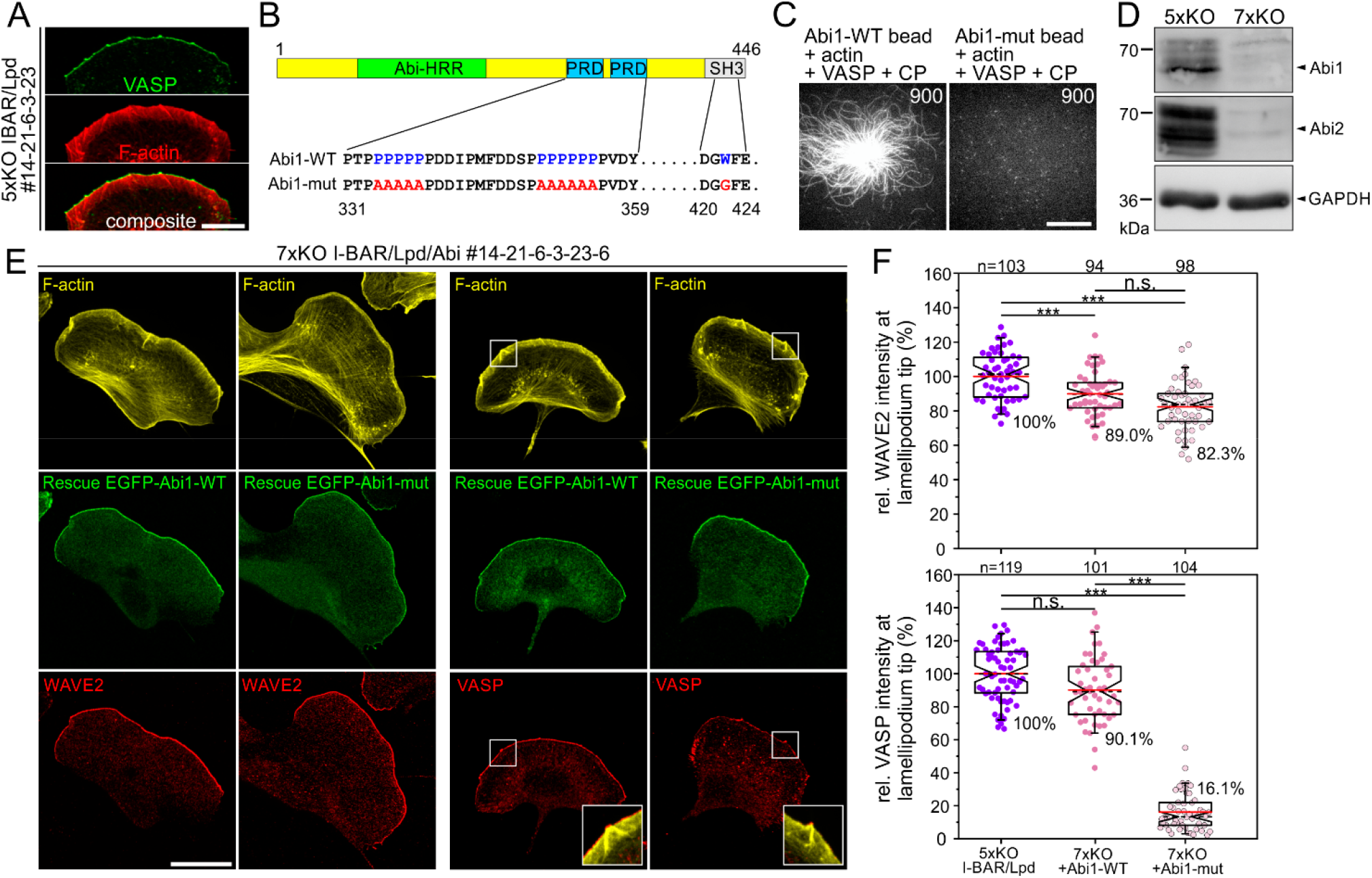
Abi1 recruits VASP to the leading edge, but is not essential for VASP clustering at microspike tips. (*A*) 5xKO cells lacking I-BAR proteins and Lpd expression still form VASP clusters at microspike tips. Gallery displays a cell stained for F-actin and endogenous VASP. Bar, 20 μm. (*B*) Domain organization of Abi1 depicting the homeodomain homologous region (HHR), the proline-rich regions (PRD) and the C-terminal SH3 domain. Numbers indicate amino acid residues. In the Abi1 mutant (Abi1-mut), the indicated proline (P) residues of the PRD were substituted with alanine (A) residues, and the highlighted tryptophan (W) was replaced by glycine (G) to inactivate the SH3 domain. (*C*) Abi1^328-481^-WT but not the Abi1-mut were able to recruit soluble VASP and initiate processive actin-filament elongation on the surface of SNAP-Abi1-derivatized beads in the presence of 1 μM G-actin (10% Atto488-labelled) and 25 nM CP. Representative images from TIRF-movies are shown. Time in sec. Bar, 10 μm. (*D*) Combined loss of Abi1 and 2 in the 7xKO I-BAR/Lpd/Abi mutant was confirmed by immunoblotting. Loading control: GAPDH. (*E*) Representative images of 7xKO mutant cells (#14-21-6-3-6) reconstituted with full-length Abi1-WT and Abi1-mut fused to EGFP. Images display cells stained for F-actin and expressed EGFP-tagged or endogenous proteins as indicated. Insets, enlarged images of boxed region. Bar, 20 μm. (*F*) Quantification of WAVE2 and VASP intensities at the tips of lamellipodia in control 5xKO and reconstituted 7xKO cells. Notched boxes in box plots indicate 50% (25-75%) and whiskers 90% (5-95%) of all measurements; arithmetic means are highlighted in red and with dashed black lines depicting the medians. WAVE2 data sets were analyzed by 1-way ANOVA and VASP data sets by Kruskal-Wallis test. ***p≤0.001; n.s.; not significant. n, number of cells. F shows pooled data from three independent experiments.

**Fig. 4.**
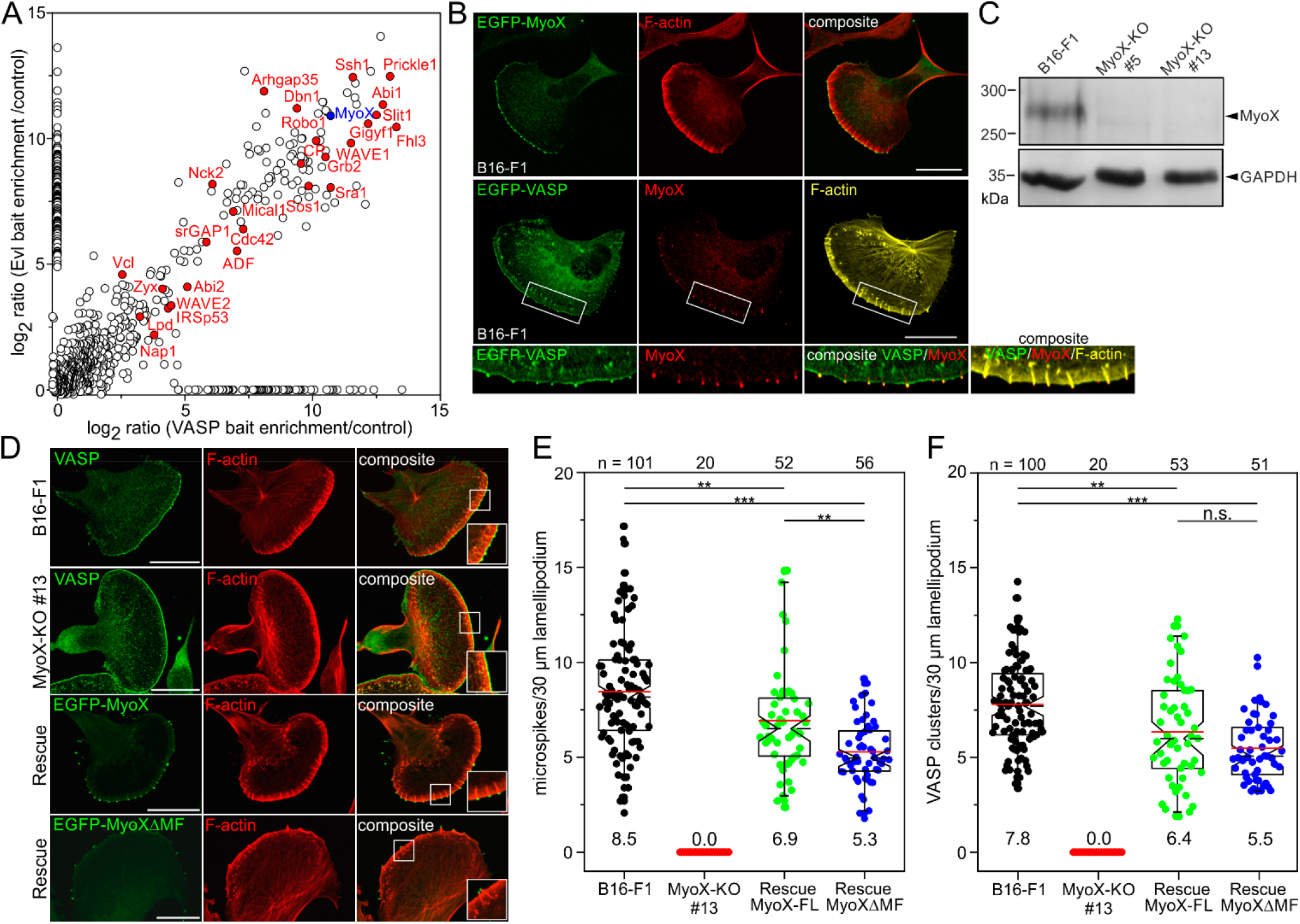
Loss of MyoX eliminates microspike formation due to abolished VASP clustering. (*A*) Graphical representation of combined MS data from BioID2 pulldowns. Enrichment of identified proteins in BioID2-Evl and -VASP pulldowns over control is shown. Proteins implicated in migration, adhesion or signaling are highlighted in red. (*B*) Ectopically expressed and endogenous MyoX accumulates at microspike tips like VASP. Cells stained with phalloidin and for endogenous or EGFP-tagged MyoX. Bars, 20 μm. (*C*) Confirmation of MyoX loss in two, independently generated MyoX-KO clones by immunoblotting. Loading control: GAPDH. (*D*) B16-F1 and MyoX-KO cells stained for endogenous VASP and F-actin showing complete loss of microspikes and VASP clusters in the mutant despite prominent localization of VASP at the leading edge (upper and middle panel). Re-expression of EGFP-MyoX and of MyoXΔMF in mutant cells rescues microspike formation (lower two panels). Bars, 20 μm. (B and D) Insets, enlarged images of boxed regions. (*E-F*) Quantification of microspikes and VASP clusters in wild-type, mutant and reconstituted cells. Notched boxes in box plots indicate 50% (25-75%) and whiskers 90% (5-95%) of all measurements; arithmetic means are highlighted in red and dashed black lines depicting the medians. (E, F) Statistical analysis by Kruskal-Wallis test. *p≤0.05; ***p≤0.001; n.s.; not significant. n, number of analyzed cells. E and F show pooled data from at least three independent experiments.

### Myosin-X is essential for VASP clustering and microspike formation

To expedite identification of proteins relevant for Ena/VASP clustering, we employed a BioID approach, which exploits a promiscuous *Aquifex aeolicus* biotin ligase (BioID2) fused to a bait that biotinylates proteins in close proximity (52). To identify exclusive or shared components in protein complexes, we generated BioID2 baits with full-length VASP and Evl, but omitted Mena due to its tendency to form aggregates. The empty BioID2 plasmid was used as negative control. After transfection of respective constructs into Ena/VASP-deficient B16-F1 mouse melanoma cells (24), protein expression in B16-F1 cells was confirmed by silver staining of analytic samples from BioID2 pulldowns on streptavidin-coated Dynabeads (SI Appendix, Fig. S13A). As opposed to empty control, both BioID-VASP and -Evl rescued microspike formation in cells, and localized to and biotinylated proteins at focal adhesions and in the leading edge including the microspike tips (SI Appendix, Fig. S13B and C). To reveal the identity of endogenous proteins, large-scale pull downs of biotinylated proteins were analyzed by mass spectrometry (MS) analysis. Besides numerous proteins implicated in actin dynamics and Ena/VASP binding including IRSp53, Lpd, and Abi, our results also interestingly revealed an almost 2000-fold enrichment of unconventional myosin-X (MyoX, Myo10) with both baits over control (Fig. 4*A*). MyoX is a widely expressed, plus-end directed molecular motor that accumulates at filopodial tips and has been implicated in VASP transport to these sites (45, 53). The knockdown of MyoX has previously been reported to decrease filopodia in Hela cells (13). Quantification of filopodia in endothelial cells derived from Myo10-null mouse strains subsequently revealed a ~50% decrease in filopodia, strongly suggesting that MyoX promotes the formation of normal filopodia numbers (54). Comparable to VASP, we identified ectopically expressed, EGFP-tagged MyoX as well as endogenous MyoX in prominent clusters at the tips of microspikes in B16-F1 cells (Fig. 4*B*). Although filopodia and microspikes represent different molecular entities (17, 24), previous work has also indicated suppression of microspikes upon siRNA-mediated knockdown of MyoX in COS7 cells (55). Thus, to determine whether this phenomenon can be extended to other cell lines, we disrupted the *Myo10* gene encoding MyoX in B16-F1 cells by CRISPR/Cas9. Loss of MyoX was confirmed by immunoblotting in independent mutants (Fig. 4*C*). Due to their highly similar phenotype, we then analyzed the morphology and F-actin distribution of B16-F1 wild-type *versus* MyoX-deficient cells (clone #13) on laminin after phalloidin staining (SI Appendix, Fig. S14A). In contrast to controls, in which approximately 60% of cells formed lamellipodia in these experiments and conditions, this fraction was increased to over 80% in MyoX-deficient cells (SI Appendix, Fig. S14B). In addition, the mutant cells frequently displayed multiple fronts compared with wild-type cells, which typically formed one leading edge (SI Appendix, Fig. S14C). Interestingly, despite a comparable F-actin content in lamellipodia, the mutant cells exhibited 15% narrower lamellipodia compared to control (SI Appendix, Figs. S14D and E). Yet, loss of MyoX resulted in significantly decreased random migration rate (30% lower) as compared to control (SI Appendix, Fig. S14F). Although this was accompanied by a 15% increase in directionality, presumably because of their much lower protrusion rates, the MyoX-KO mutants also showed lower MSD values as compared to B16-F1 control (SI Appendix, Figs. S14F-I). This was confirmed by analyses in the independent MyoX-KO clone #5 (SI Appendix, Fig. S15). Notwithstanding this, the distribution and intensity of the prominent lamellipodia markers VASP, WAVE2 and cortactin in MyoX-null cells remained virtually unchanged compared to wild-type (SI Appendix, Fig. S16).

Next, we analyzed microspike formation and VASP clustering in wild-type, mutant and reconstituted cells. In contrast to B16-F1 cells, which displayed prominent microspikes embedded into the protruding cell edge, MyoX-null cells exhibited a complete loss of microspikes associated with a total absence of VASP clusters (Figs. 4*D-F* and Movie S6). Given the evident lack of microspikes, we stained for the actin-crosslinking protein fascin, a well-established constituent of microspikes in B16-F1 cells (56). Consistently, the MyoX mutant lacked fascin-containing bundles, as opposed to wild-type, despite comparable, global expression of fascin in the mutant (SI Appendix, Fig. S17). This was corroborated by live cell imaging of wild-type and MyoX-KO cells expressing EGFP-fascin (Movie S7). To ensure that the observed phenotype was MyoX-specific, the same parameters were analyzed in independent MyoX-KO mutant cells ectopically expressing low levels of EGFP-MyoX. Re-expression of MyoX and of a MyoX variant lacking its C-terminal MyTH/FERM cargo domain (MyoXΔMF), but not a motor dead mutant (G437A) efficiently rescued microspike formation and VASP clustering at their tips (Figs. 4*D-F* and SI Appendix, Fig. S18), demonstrating not only that the MyTH/FERM is dispensable for transport of VASP to microspike tips, but also revealing a crucial role for MyoX motor activity in microspike formation. Notably, and consistent with a recent report analyzing functional domains of MyoX in filopodium formation (57), reconstituted MyoX-KO cells expressing MyoXΔMF formed shorter microspikes (20% reduction) lacking Lpd at their tips (SI Appendix, Fig. S19), once again illustrating the accessory role of Lpd in this process. Finally, we also completely abolished microspike formation and VASP clustering by eliminating MyoX in Rat2 cells by CRISPR/Cas9 (SI Appendix, Fig. S20), establishing MyoX, besides its critical function in filopodia formation, as an essential factor in the formation of microspikes and VASP clustering at their tips in different cell types.

### VASP is essential for tethering of MyoX clusters to the actin network

The impaired microspike formation in Lpd-KO mutants, their marked decrease in Ena/VASP-deficient EVM-KO cells, and their complete loss in MyoX-KO cell lines prompted us to investigate the potential interplay between these proteins in more detail. Previous work proposed initial formation of small leading-edge Lpd clusters that expand, fuse and subsequently recruit VASP to induce filopodia/microspike formation (26). We therefore examined the pairwise localization of MyoX, Lpd and VASP in all combinations in a 1 μm wide region of interest (ROI) at the leading edge in reconstituted MyoX-KO and EVM-KO cells expressing low levels of EGFP-tagged MyoX and VASP by confocal microscopy. Subsequent analyses strongly suggest colocalization of MyoX with VASP, of MyoX with Lpd and of VASP with Lpd as evidenced by respective Pearson correlation coefficients yielding values of 0.79, 0.77 and 0.83 (Figs. 5*A-C*). However, the existence of Lpd precursor assemblies devoid of VASP could not be experimentally confirmed (Fig. 5*C*). Given the striking correlation coefficient of 0.84, MyoX and VASP also appear to perfectly colocalize in Lpd-deficient cells (Fig. 5*D*), strongly suggesting that Lpd does not operate upstream of MyoX in the molecular pathway leading to Ena/VASP clustering. In line with this notion and as opposed to wild-type or EVM-KO cells, MyoX-deficient cells were incapable of forming any Lpd clusters (Figs. 5*E* and *F*). Thus, due to the outstanding role of MyoX and VASP in this process, we used TIRF time-lapse microscopy to compare the behavior of MyoX clusters in MyoX-KO and EVM-KO cells expressing low levels of EGFP-MyoX. In stark contrast to reconstituted MyoX-KO cells, in which the MyoX clusters translocated predominantly straight forward within the advancing cell front largely as expected from known microspike behavior (7), we observed a dramatically increased lateral movement of MyoX clusters along the lamellipodium tip in EVM-KO cells (Fig. 5*G* and Movie S8). This was corroborated by comparison of MyoX cluster dynamics in reconstituted MyoX-KO *versus* EVM-KO cells by adaptive kymograph analyses, in which the intensity profiles of MyoX in the advancing leading edge were plotted over time (Fig. 5*H*). Quantifications revealed that the lateral speed of EGFP-MyoX clusters in the EVM-MO mutant (with 5.15±0.99 μm min^-1^) was on average nearly one order of magnitude faster than that in reconstituted MyoX-KO cells (0.61±0.51 μm min^-1^), and still about 4 times faster than the fastest laterally moving MyoX clusters in reconstituted MyoX-KO cells (Fig. 5*I*). This implies that in the absence of Ena/VASP, MyoX clusters, which also contain Lpd (SI Appendix, Fig. S21), are not accurately tethered to the lamellipodial actin network. To test this, we examined F-actin accumulation beneath MyoX clusters in both cell lines (Fig. 5*J*). In contrast to reconstituted MyoX-KO cells where the clusters localized at the tips of microspikes that were oriented close to perpendicular to the membrane, in EVM-KO cells we detected short, tortuous actin comets at the MyoX clusters that nestled against the membrane. Notably, these comets did not contain fascin, as opposed to the microspikes of reconstituted MyoX-KO cells (Fig. 5*J* and Movie S9). Combined, these findings not only demonstrate that MyoX is obligatory for clustering of Ena/VASP proteins to initiate processive actin-filament elongation at microspike tips. They further imply that the development of genuine microspikes initiated by a functional tip complex harboring MyoX and Lpd specifically requires Ena/VASP proteins, for tethering these complexes to dendritic precursor filaments to promote processive actin assembly of bundles stabilized by shaft proteins including fascin.

**Fig. 5.**
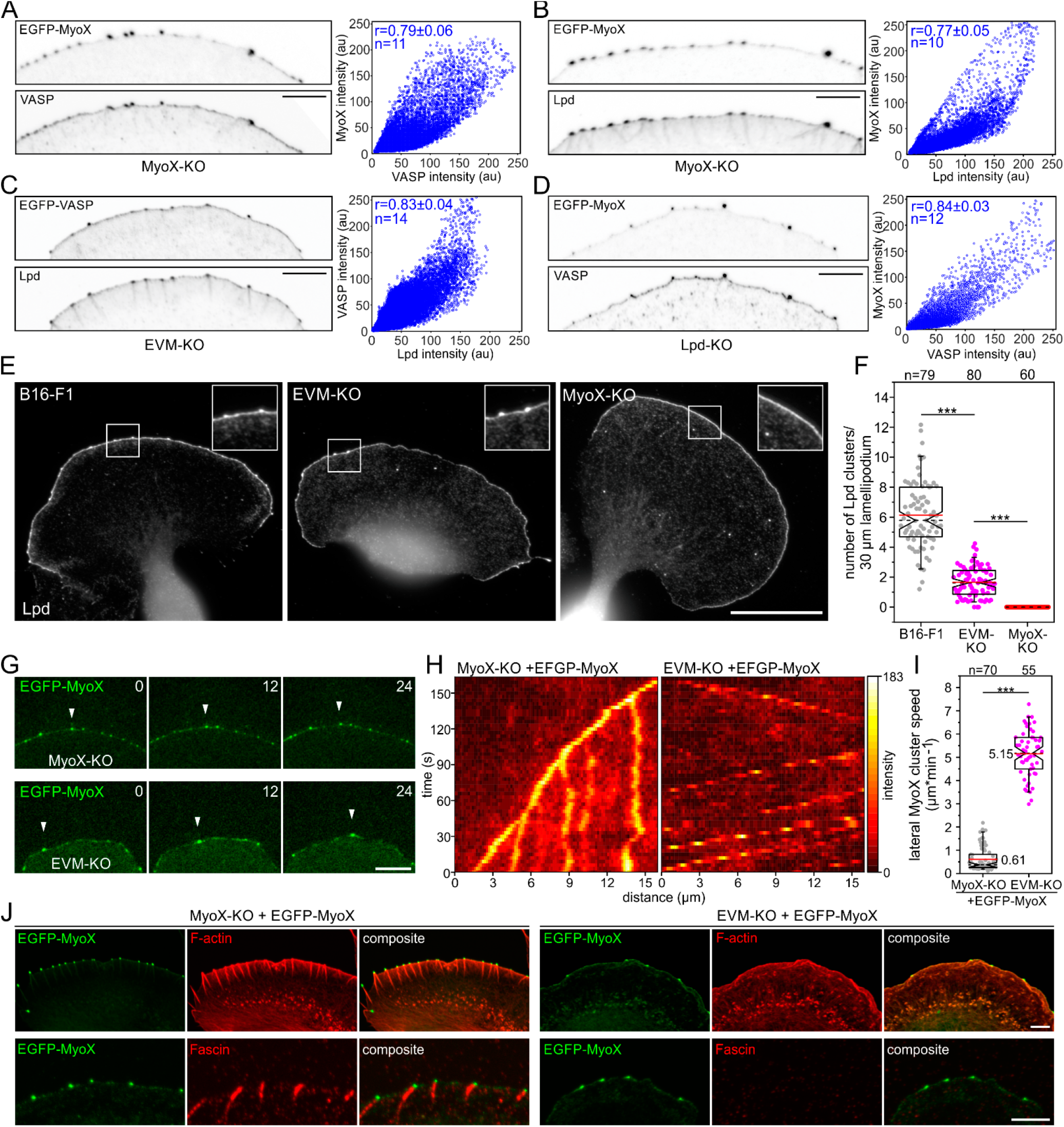
Loss of Ena/VASP dramatically increases lateral flow of MyoX clusters at the leading edge. (*A-C*) VASP, MyoX and Lpd colocalize in microspike tip clusters. Inverted images of representative leading edges in reconstituted MyoX-KO and EVM-KO cells expressing constructs as indicated (*left panels*). Bar, 10 μm. The intensities of respective protein pairs were measured in an 1 μm wide sector of the leading edge to generate cytofluorogram with respective intensity distributions (*right panels*) and to calculate corresponding Pearson correlation coefficients (r) and mean±SD. (*D*) Equivalent analysis of MyoX and VASP in Lpd-deficient cells. (*E*) Loss of MyoX completely suppressed Lpd cluster formation, whereas EVM-KO cells were still able to form some Lpd clusters. Representative images of wild-type, EVM-KO and MyoX-KO cells stained for endogenous Lpd. Insets, enlarged images of boxed regions. Bar, 20 μm. (*F*) Quantification of Lpd clusters in wild-type and mutant cells. (*G*) Representative images from TIRF microscopy time-lapse movies of MyoX-KO or EVM-KO mutant cells expressing low levels of EGFP-MyoX. White arrowheads indicate the position of the same MyoX cluster in respective cell lines over time. Bar, 10 μm. (*H*) Adaptive kymographs illustrate dramatically increased lateral flow of MyoX clusters in EVM-KO cells as opposed to reconstituted MyoX-KO cells. Heat map indicates pixel intensities in 8-bit unsigned integer range. (*I*) Quantification of lateral MyoX cluster speed at the leading edge in MyoX-KO or EVM-KO mutant cells expressing low levels of EGFP-MyoX. (*J*) Representative images of MyoX-KO and EVM-KO cells expressing EGFP-MyoX and stained for Factin (*upper panel*) or endogenous fascin (*lower panel*). Bars, 5 μm. Notched boxes in the box plot indicate 50% (25-75%) and whiskers 90% (5-95%) of all measurements; arithmetic means are highlighted in red and dashed black lines depicting the medians. (F, I) Non-parametric, Mann-Whitney-U-test was used for statistical analysis. ***p≤0.001. n, number of cells (F) or clusters (I). A-D, F and I show pooled data from three independent experiments.

## Discussion

In this work, we dissected the molecular pathways required for VASP clustering and microspike formation in B16-F1 cells. IRSp53 was a prime candidate since it localizes to lamellipodia and filopodia tips and has been assumed an essential regulator of filopodia formation (30). Moreover, it is capable of recruiting and clustering VASP on beads or synthetic membranes to initiate processive actin assembly *in vitro* (25, 27). Consistent with the ability of I-BAR proteins to directly interact with Ena/VASP proteins (30), the 4xKO mutants exhibited diminished VASP recruitment to the leading edge associated with reduced cell migration rates. Unexpectedly, however, in the cellular context we found no evidence for a critical role of IRSp53 or one of the other I-BAR family proteins in VASP clustering or microspike formation. Thus, the previously reported VASP recruitment in *in vitro* experiments could just lead to molecular crowding of VASP on the bead surface, which is dense enough to mimic clustering and initiate actin assembly. To our surprise, the loss of I-BAR proteins did also not decrease the formation of filopodia induced by expression of active mDia2 or MyoX. The combined loss of the most abundant I-BAR family members IRSp53 and IRTKS in NIH 3T3 fibroblast confirmed this view, which overall argues against a prominent role for these proteins in filopodia formation. Nevertheless, I-BAR protein deficiency resulted in quite chaotic cell morphologies, with much lower fractions of cells forming smooth lamellipodia. This might be either caused by reduced levels of Ena/VASP proteins and the Arp2/3 complex activator WAVE2 at the cell front, the diminished global levels of the WRC components WAVE2 and Abi or perturbation of other signaling pathways that are known to coordinate the actin cytoskeleton (58).

Consistent with the presence of 6 EVH1-binding sites for Ena/VASP proteins in Lpd (26, 34), we found its loss to be associated with reduced recruitment of VASP to the leading edge in the 1xKO mutant. This was particularly relevant in the context of reduced microspike formation in the absence of Lpd, a parameter not explored in our initial characterizations of Lpd-deficient cells (35). The accompanying, moderate decrease in lamellipodial F-actin intensity and width also teased out for Lpd single-KO cells in current analyses, perfectly matches all these observations. Notably, the latter has also been missed in our previously published experiments (35), partially because of being subtle, more likely though due to potential variations in reagents, fixation protocols and experimental conditions. Notwithstanding this, all these new data are fully compatible with our previous conclusion that Lpd is not essential for, but optimizes cell edge protrusion (35). Unexpectedly, however, the combined loss of I-BAR family members and Lpd resulted in a reversion of the 4xKO phenotype, both with respect to recruitment of VASP or other relevant lamellipodial factors and to F-actin distribution, although these cells also exhibited reduced expression levels of the WRC components WAVE2 and Abi. The precise mechanistic reasons for these observations, however, have remained unclear and need to be explored in future work. Consistent with previous work using the *Drosophila* (48) or *Dictyostelium* model systems (47), and as also suggested by the analysis of B16-F1 cells lacking functional WRC, but harboring non-canonical, lamellipodia-like structures (51), we find here that the WRC subunit Abi plays a major role in the recruitment of VASP to the leading edge in mammalian cells. In contrast to the loss of Lpd, however, the combined loss of Abi1 and 2 on top of I-BAR and Lpd (7xKO) did not noticeably perturb VASP clustering at the cell edge.

Mechanistically, the actin filament-binding activity of Lpd, which can occur independently of Ena/VASP binding, was proposed to elicit Ena/VASP clustering and tether the complexes to growing barbed ends, thereby boosting their processive polymerase activity (34). However, VASP-mediated actin assembly on beads conjugated with the actin-binding deficient mutant Lpd-44A was even stronger as compared to Lpd-WT in the presence of CP (Fig. 2*B*). Moreover, filaments on Lpd-44A beads also grew about three times faster thus forming longer filaments as compared to Lpd-WT control, clearly arguing against decreased VASP processivity in the presence of the Lpd-44A mutant. Most importantly, expression of Lpd-44A was effective in rescuing the microspike defect of 5xKO I-BAR/Lpd mutants in a fashion comparable to full length Lpd-WT, in spite of the failure of the mutant to target to the leading edge. Thus, in cells the polybasic clusters located within the putative, C-terminal actin-binding site of Lpd could regulate its membrane positioning and/or stabilization by interaction with acidic phospholipids, as shown previously for IRSp53 (59). However, *in vitro* and in the absence of these phospholipids, Lpd-WT is likely to compete with VASP for recruitment of the negatively charged actin monomers, which in turn is expected to reduce overall charging of VASP with monomers, and hence, the VASP-mediated filament elongation (5). For the reasons stated above, and given that RIAM, the second member of the two-protein MRL family in vertebrates is also absent in B16-F1 cells (35), the mechanism proposed for Lpd family mode of action appears unlikely. On the other hand, Lpd does clearly contribute to microspike formation, as average microspike numbers are reduced to half upon Lpd removal. Moreover, F-actin intensities in microspikes of Lpd-mutants are reduced by ~20%, suggesting fewer filaments in these bundles. Thus, although not essential, Lpd constitutes a relevant accessory protein supporting Ena/VASP clustering and promoting the formation of microspikes with optimal filament numbers.

Here, we also identified MyoX as factor proximal to VASP and Evl, and showed that besides its critical function in filopodia formation, MyoX is obligatory for both Ena/VASP clustering and microspike formation in different cell types. Notably, and despite normal recruitment of VASP and other relevant proteins to the edge of the lamellipodium, the latter is significantly reduced in width in MyoX-KO as compared to B16-F1 control, potentially suggesting that microspikes could support lamellipodium architecture and boost cell migration.

Our Ena/VASP BioID2 biotinylation approach allows the identification of proteins located within a ~10 nm-range to baits, so cannot discriminate between direct and indirect binding. However, MyoX and VASP were previously reported to co-immunoprecipitate (60, 61), supporting the view that both proteins are components of the same complex or even directly interact with each other. The coincident dependence of MyoX and Ena/VASP proteins on microspike formation is reminiscent of equivalent observations for DdMyo7 and DdVASP in filopodia formation in *Dictyostelium*, where loss of either protein abrogates the process (20, 62). Nevertheless, there are some marked differences, for instance regarding recruitment mechanisms. In *Dictyostelium*, DdMyo7 fails to be recruited to the leading edge in the absence of DdVASP, implying that DdVASP-mediated actin assembly and bundling is needed for DdMyo7 activation (63). In contrast, MyoX acts upstream of Ena/VASP and Lpd in microspike formation in B16-F1 cells, since even after Ena/VASP removal or lack of Lpd recruitment in MyoX-KO cells expressing MyoXΔMF, MyoX clusters at the leading edge are still being formed. Moreover, while expression of a constitutively active filopodial formin partly rescued DdMyo7 recruitment and filopodium formation of DdVASP-deficient cells (63), formin-driven actin assembly by active variants of mDia2, FMNL2 or -3 were all insufficient to rescue the defect in microspike formation in Ena/VASP-deficient B16-F1 cells (24).

Notably, MyoX has been previously implicated in intrafilopodial transport of Ena/VASP proteins towards the tip of the growing filopodium (60). Although this would be mechanistically sufficient for explaining sustained elongation of filopodia at steady state, it falls short in explaining the formation of a functional tip complex to initiate filopodium formation. Assuming that clustering of Ena/VASP proteins is instrumental for initiating processive actin assembly in the presence of CP, we can sketch a working model for microspike formation in mammalian cells. The MyoX tail encompasses a coiled-coil region for antiparallel dimerization and three pleckstrin homology (PH) domains that target MyoX to phosphatidylinositol (3,4,5)-trisphosphate (PIP3)-rich regions of the plasma membrane, followed by a MyTH4-FERM (MF; myosin tail homology - band 4.1, ezrin, radixin, moesin) domain at its C-terminus (64). MyoX is assumed to be a monomer prior to activation, but binding to PIP3 apparently opens up the molecule and promotes activation and dimerization at the leading edge. By hitherto uncharacterized activities, MyoX dimers then associate to higher-order clusters, which also harbour Lpd. These clusters in turn are likely to recruit and cluster Ena/VASP proteins and tether them onto lamellipodial actin network filaments. The accompanied, processive assembly of parallel actin filaments drives the emergence of bundles, which are continuously stabilized by fascin to yield microspikes (56).

## Material and Methods

### Constructs

Constructs used in this study: pEGFP-C1-IRSp53, pEGFP-C1-IRTKS, pEGFP-C1-ABBA, pEGFP-C1-MIM, pEGFP-C1-Abi1, pEGFP-C1-Abi1-mut, pEGFP-Lpd (33), pEGFP-C1-Lpd44A, Lifeact-EGFP (65), pEGFP-C1-MyoX, pEGFP-C1-MyoX(G437A), pEGFP-C1-MyoXΔMF, pEGFP-C1-mDia2ΔDAD (12), pEGFP-C1-VASP (24), pmCherry-VASP, pmCherry-MyoX, pGEX-6P1-IRSp53, pGEX-6P1-IRTKS, pGEX-6P1-Pinkbar, pGEX-6P1-ABBA, pGEX-6P1-MIM, pGEX-6P1-hVASPdDGAB4M, pQE-30-SNAP-GCN4-Lpd^850-1250^WT, pQE-30-SNAP-GCN4-Lpd^850-1250^44A, pQE-30-SNAP-GCN4-Abi1^328-481^-WT, pQE-30-SNAP-GCN4-Abi1^328-481^-mut, pQE-30-SNAP-GCN4-Abi2^326-446^-mut, pQE-30-MyoX^1197-1509^, pQE-30-CapZ (66). Detailed information is described in SI Appendix, Materials and Methods.

### Semi quantitative RT-PCR

Total RNA of B16-F1 cells by standard procedures. Respective cDNAs were synthesized with the Maxima H Minus First Strand cDNA Synthesis Kit. Subsequently, respective control plasmids and B16-F1 cDNA were used as templates for PCR reactions to assess relative I-BAR protein expression levels in B16-F1 cells. Detailed information is described in SI Appendix, Materials and Methods.

### Cell culture, transfection, and genome editing by CRISPR/Cas9

sgRNAs for elimination of mouse *Irsp53, Irtks, Mtss2, Mtss2, Abi1, Abi2* and mouse or rat *Myo10* were ligated into expression plasmid pSpCas9(BB)-2A-Puro(PX459)V2.0 (67). The targeting construct for the *Raph1* gene has been reported (35). After transfection and selection with puromycin of B16-F1, NIH 3T3, Rat2 and derived cells, routinely cultivated at 37°C and 5% CO_2_ in high-glucose DMEM culture medium supplemented with 1% penicillin-streptomycin, 10% FBS, and 2 mM UltraGlutamine, clonal cell lines were validated by the TIDE sequence trace decomposition web tool (40) and immunoblotting. Detailed information is described in SI Appendix, Materials and Methods.

### Protein purification

GST- and His-tagged fusion proteins were expressed in *E. coli* strains Rossetta 2 or M15, respectively, and purified by affinity chromatography according to the instructions of the manufacturers. The tags were cleaved off by PreScission protease (GE Healthcare) and the proteins were further purified by size-exclusion chromatography (SEC) using an Äkta Purifier System. The purification of capping protein (66) and actin (68) have been described. Detailed information is described in SI Appendix, Materials and Methods.

### Immunoblotting and immunofluorescence

Immunoblotting and immunofluorescence were performed essentially as described (24). Detailed information is described in SI Appendix, Materials and Methods.

### Live cell imaging

Live cell imaging was performed essentially as described (24). Detailed information is described in SI Appendix, Materials and Methods.

### Induction of filopodia in absence of lamellipodia

Induction of filopodia was performed essentially as described (24). Detailed information is described in SI Appendix, Materials and Methods.

### TIRF assays

For TIRF bead assays SNAP-capture magnetic beads were coated with 10 μM of SNAP-GCN4-Lpd^850-1250^, -Lpd^850-1250^(44A), -Abi1^331-424^-WT or -Abi1^331-424^-mut and were subsequently split into two fractions. One was incubated with coupling buffer and served as negative control, while the other was incubated with 0.5 μM soluble His-VASPDdGAB in coupling buffer. TIRF assays were performed as previously described (18). Images were acquired with a Nikon Eclipse TI-E inverted microscope equipped with a TIRF Apo 100x objective. Detailed information is described in SI Appendix, Materials and Methods.

### BioID2 pulldown and mass spectrometry

BioID2 assays and mass spectrometry were performed essentially as previously reported (52, 69). Detailed information is described in SI Appendix, Materials and Methods.

### Statistical analyses

Statistical analyses was performed as described (24). Detailed information is described in SI Appendix, Materials and Methods.

## Acknowledgements

We thank Brigitte Jockusch and Sabine Buchmeier for fascin antibody. This work was supported in part by the Deutsche Forschungsgemeinschaft (individual grants FA330/9-2 and FA330/13-1 to J.F. and Research Training Group GRK2223/1 to K.R.), and by funding from the Helmholtz Society (to K.R.). GS was supported by the Associazione Italiana per la Ricerca sul Cancro (AIRC IG#18621 and 5Xmille#22759) and the Italian Ministry of University and Scientific Research (PRIN 2017, 2017HWTP2K).

