## Supplemental Information for "Ena/VASP clustering at microspike tips involves Lamellipodin but not I-BAR proteins, and absolutely requires unconventional Myosin-X"

**Supplementary Information**

### Materials and Methods

#### Constructs

For expression of pEGFP-tagged proteins, full-length cDNAs encoding IRSp53, IRTKS, ABBA, MIM, Abi1 and Abi2 were amplified as BamHI-SalI fragments from B16-F1 cDNA and inserted into the BglII-SalI sites of pEGFP-C1 (Clontech). pEGFP-C1-Lpd has been described (1). After partial digest of pEGFP-C1-Lpd with AgeI and blunting of this site located upstream of EGFP, pEGFP-C1-Lpd44A was generated by cleavage of pEGFP-C1-Lpd(ΔAgeI) with AgeI and SacII and replacement of the corresponding region obtained from pUC-Lpd$^{850\text{-}1250}$44A. EGFP-C1-Abi1-mut was generated by cleavage of pEGFP-C1-Abi1 with BspEI and XhoI and replacement of the corresponding region obtained from pUC-Abi1-mut that has been synthesized (BioCat), and in which the proline residues 334-338 and 350-355 were mutated to alanine residues. It additionally harbors point mutation W421G in the SH3 domain to prevent interactions with proline-rich regions of other proteins. Full-length mouse MyoX was amplified from B16-F1 cDNA and ligated into the BglII and Asp718I sites of pEGFP-C1 and pmCherry (Addgene ID: 632524). Murine MyoX lacking the C-terminal MyTH/FERM domain (MyoXΔMF, aa 1-1509), was amplified from pEGFP-C1-MyoX and ligated into the BglII and SalI sites of pEGFP-C1. The MyoX motor-dead mutation (G437A) was generated by side-directed mutagenesis using pEGFP-C1-MyoX as template, as described previously for bovine MyoX (2). Lifeact-EGFP (3), pmCherry-VASP, pEGFP-C1-VASP (4) and EGFP-C1-mDia2ΔDAD (5) have been described. For the generation of BioD2 baits, VASP and Evl were amplified from respective pEGFP-C1 plasmids (4) and ligated into XhoI and EcoRI sites (VASP) or XhoI and BamHI sites (Evl) of myc-BioID2-MCS (Addgene ID: 74223, (6)).

For protein purification and PCR analyses, fragments encoding full-length IRTKS, IRSp53, ABBA and MIM were amplified from respective pEGFP-C1 plasmids and inserted into the same sites BamHI-SalI of pGEX-6P1 (GE Healthcare). Full-length Pinkbar was synthetized (BioCat) and inserted into the BamHI-SalI sites of pGEX-6P1. Human VASP harboring the high affinity G-actin binding site from *Dictyostelium* (7), was excised with BamHI and SalI from pGEX-6P1-hVASPdDGAB4M (8) and inserted into the same sites of pQE-30 (Qiagen). For TIRF experiments using SNAP-GCN4-tagged fusion proteins plasmid pQE-30-P-SNAP-GCN4 was designed. A BglII-BamHI fragment encoding the PreScission protease cleavage site (P) followed by the SNAP-tag was amplified from pGEX-6P1-SNAP (8) and inserted into the BamHI site of pQE-30, yielding pQE-30-P-SNAP. Subsequently, a BglII-BamHI fragment encoding the GCN4 dimerization coiled-coil domain was amplified from pGEX-6P1-hVASPdDGAB2M (8) and inserted into the BamHI site of pQE-30-P-SNAP yielding pQE30-P-SNAP-GCN4. Subsequently, a fragment encoding the C-terminus of human Lpd (Lpd$^{850-1250}$) was amplified from pEGFP-C1-Lpd (1) and inserted into the same sites of pQE30-P-SNAP-GCN4. Mutant Lpd$^{850-1250}$44A (9), in which all basic residues were replaced with alanine residues, was synthetized (BioCat) with BamHI-SalI restriction sites and generated accordingly. The C-terminal regions of mouse Abi1 and Abi1-mut (328-481) and Abi2 (326-446) were amplified from B16-F1 cDNA and inserted into the BamHI-SalI site of pQE30-P-SNAP-GCN4. For generation of antibodies, a MyoX fragment encoding residues 1197-1509 was amplified from pEGFP-C1-MyoX, and inserted into the BamHI and SalI sites of pQE-30. The plasmid for expression of His-tagged CapZ has been reported (7). All generated constructs were validated by sequencing.

**Semi quantitative RT-PCR**

Total RNA of B16-F1 cells was isolated with the RNAeasy kit (Qiagen). Respective cDNAs were synthesized with the Maxima H Minus First Strand cDNA Synthesis Kit (Thermo Scientific) using 10 µg of total RNA as template. For PCR reactions, the PCR primers were designed to span exon-exon boundaries, precluding amplification of genomic fragments. Efficacy of primer pairs was validated by conventional PCR using 50 ng of pGEX-6P1 constructs harboring cDNAs encoding full-length IRSp53, IRTKS, Pinkbar, ABBA, and MIM, respectively. Subsequently, 50 ng of cDNA was used as template in PCR reactions to assess relative I-BAR protein expression levels in B16-F1 cells.

**Cell culture and transfection**

B16-F1 mouse melanoma (ATCC CRL-6323), NIH 3T3 (ATCC CRL-1658) and Rat2 (ATCC CRL-1764) fibroblasts as well as derived mutant cell lines were cultured at 37°C and 5% $CO_2$ in high-glucose DMEM culture medium (Lonza) supplemented with 1% penicillin-streptomycin (Biowest), 10% FBS (Biowest), and 2 mM UltraGlutamine (Lonza). After seeding onto 35 mm diameter wells (Sarstedt), cells were transfected with 1 µg (B16-F1 and Rat2 cells) or 3 µg (NIH 3T3 cells) plasmid DNA using JetPRIME transfection reagent (PolyPlus) according to the manufacturer's protocol. Used cell lines were routinely authenticated following common guide lines by local authorities.

**Genome editing by CRISPR/Cas9**

DNA target sequences of respective genes were pasted in the CRISPR/Cas9 design tool (https://cctop.cos.uni-heidelberg.de) to generate sgRNAs of 20 nucleotides with high-efficiency scores and minimal off-target efficiency covering all possible splice variants. For inactivation of the *Irsp53* gene the sgRNA 5'-CCATGGCGATGAAGTTGCGG-3' targeting exon 2, for inactivation of the *Irtks* gene the sgRNA 5'-AAAAGCCTACTACGACGGCG-3' (exon 3), for inactivation of the *Mtss2* gene, the sgRNA 5'-TGCAGGAGCGCATCGAAGAC-3' (exon 3) and for inactivation of the *Mtss1* gene, the sgRNA 5'-TTTCTGAAAGGCGTCCAAGA-3' (exon 3) were used. For disruption of the *Abi1* gene, the sgRNA 5'-AGGAGATCCCGTCTGGCAAG-3' (exon 1), for *Abi2,* the sgRNA 5'-TGAGTTGGAGGTGTCGTACG-3' (exon 4), for mouse *Myo10,* the sgRNA 5'-GTGATCCGAGCTCACATCTT-3' (exon 22) and for rat *Myo10* sgRNA 5'- ATGTGAGCTCGGATGACCA-3' (exon 22) were used. All sgRNAs were ligated into expression plasmid pSpCas9(BB)-2A-Puro(PX459)V2.0 (Addgene plasmid ID: 62988) using BbsI (10) and sequences validated by sequencing. The targeting construct for the *Raph1* gene has been reported (11). 24 h after transfection, the cells were selected for 4 days with culture medium containing 3 $\mu g \cdot ml^{-1}$ puromycin (Invivogen) and then cultivated for 24 h in the absence of puromycin. Single cells were seeded into 96-well microtiter plates by visual inspection and expanded for isolation of clonal cell lines in pre-conditioned medium without puromycin. Subsequently, suitable fragments of about 500 bp spanning the respective target sites were amplified from genomic DNA, and clones analyzed by the TIDE sequence trace decomposition web tool (http://shinyapps.datacurators.nl/tide/; (12)).

**Purification of recombinant proteins**

For expression, the GST-fusion proteins constructs were transformed into *E. coli* strain Rossetta 2 (Novagen) and protein expression induced with 0.75 μM isopropyl-β-D-thiogalactoside (IPTG, CarlRoth) at 21 °C for 14 h. After harvesting, the cells were resuspended in lysis buffer containing 30 mM HEPES, pH 7.4, 300 mM NaCl, 2 mM EDTA, 2 mM benzamidine, 1 mM DTT, 5% (vol/vol) glycerol, 0.1 mM AEBSF (AppliChem) and 2 units·$ml^{-1}$ benzonase (Merck-Millipore) and lysed by ultrasonication. Fusion proteins were subsequently purified from bacterial extracts by affinity chromatography using Protino Glutathione Agarose 4B (Macherey-Nagel) using standard procedures. After cleaving off the GST-tag with PreScission protease (GE Healthcare), the proteins were further purified by size-exclusion chromatography (SEC) using HiLoad 26/75 Superdex, HiLoad 26/600 Superdex 200 or Superose 6 columns (GE Healthcare) controlled by an Äkta Purifier System. Constructs encoding His-Tag fusion proteins were transformed into the host strain *E.coli* M15 (Qiagen). Purification of His-tagged proteins with cOmplete His-Tag Purification Resin (Sigma) was performed according to the instruction of the manufacturer. The murine MyoX fragment encoding the three PH-domains (aa1197-1509) was expressed as His-tag fusion protein and purified by affinity chromatography using cOmplete His-Tag Purification Resin under denaturing conditions according to the manufacturer's instructions. The His-tag of SNAP-GCN4 fusion proteins used for TIRF experiments was cleaved off with PreScission protease and the proteins further purified by SEC. The purification of human heterodimeric capping protein (CapZ) has been be described (7). Purified proteins were stored at -20 °C in storage buffer (50 mM HEPES, pH 7.4, 150 mM NaCl, 1 mM DTT and 60% (vol/vol) glycerol). Actin was extracted and purified from acetone powder of rabbit skeletal muscle following standard procedures. Fractions were labelled on Cys374 with Atto488-maleimide (ATTO-TEC) and stored in G-Buffer (5 mM Tris/HCl pH 8, 0.2 mM ATP, 0.5 mM DTT, 0.2 mM $CaCl_2$, 0.1 $mg \cdot ml^{-1}$ $NaN_3$).

**Antibodies used**

Polyclonal antibodies directed against IRSp53, IRTKS, MyoX, Abi 1 and 2 were raised by immunizing female New Zealand white rabbits with respective recombinant proteins following standard procedures. For immunoblotting, polyclonal antibodies directed against IRSp53 (1:1000 ), IRTKS (1:1000), Lpd (1:1000), Abi1 (1:1000), Abi2 (1:1000), MyoX (1:1000), VASP (1:1000, (4)), WAVE2 (1:1000,(4)) or mouse monoclonal antibodies against CP α-subunit (undiluted hybridoma supernatant of clone mAb 5B12.3, DSHB), actin (1:1000, pan-actin-antibody [4A4], # ab119952, Abcam, Myc (1:200, clone 9E10, #sc-40, Santa Cruz Biotechnology), glyceraldehyde-3-phosphate dehydrogenase (GAPDH) (1:10,000; #CB1001-500UG, Merck-Millipore), Arp3 (1:1000; #A5979-200UL, Sigma) or fascin (undiluted hybridoma supernatant of clone 5E2) were used.
Primary antibodies in immunoblots were visualized using phosphatase-coupled anti-rabbit antibodies (1:1000; #115-055-144, Dianova) or anti-mouse (1:1000; #115-055-62, Dianova). For immunofluorescence, the following primary antibodies were used: rabbit anti-cortactin (1:1000, (4)), rabbit anti-VASP (1:1000, (4)), rabbit anti-WAVE2 (1:1000, (4)), rabbit anti-MyoX (1:1000, this study), mouse anti-Arp2/3 and mouse anti-fascin antibody 5E2 (undiluted hybridoma supernatant, (4)). Primary antibodies were visualized in immunohistochemistry with Alexa-555-conjugated goat-anti-rabbit (1:1000; #A21429,

Invitrogen), Alexa-488-conjugated goat-anti-rabbit (1:1000; #A-11034, Invitrogen), Alexa-555-conjugated goat-anti-mouse (1:1000; #A32727; Invitrogen) or Alexa-488-conjugated goat-anti-mouse (1:1000; #A11026; Invitrogen). The EGFP-signal of transfected cells expressing EGFP-fusion proteins was enhanced with Alexa488-conjugated nanobodies (4). For visualization of F-actin, Atto550-phalloidin (1:250, #AD550-82, ATTO-TEC) or Atto633-phalloidin (1:250, #AD633-81, ATTO-TEC) were used. Biotinylated proteins were visualized with Atto488-streptavidin (1:1000, #AD 488-61, ATTO-TEC).

**Immunoblotting**

For preparation of total cell lysates, cells were cultured to confluency and trypsinized. Cell pellets were washed twice with cold PBS and resuspended in ice-cold RIPA buffer (50 mM Tris/HCl, pH 8.0, 150 mM NaCl, 0.5% sodium deoxycholate, 1.0% Triton X-100, 0.1% sodium dodecyl sulfate) supplemented with 5 mM benzamidine, 0.1 mM AEBSF and benzonase (1:500) for 1 hr at 4°C on a wheel rotator. Subsequently, lysates were clarified by centrifugation for 5 min at 16,000xg and the supernatant mixed with the same amount of 3xSDS loading buffer. After SDS-PAGE the proteins were transferred onto nitrocellulose membranes (Hypermol), which were blocked with NCP buffer (10 mM Tris/HCl, pH 8.0, 150 mM NaCl, 0.02% $NaN_3$, 0.05% Tween-20) containing 4% bovine serum albumin (BSA), and incubated with primary antibodies in NCP overnight. After washing and incubation with secondary, phosphatase-conjugated antibodies, membranes were developed with 20 mg $ml^{-1}$ of 5-brom-4-chlor-3-indolylphosphat-p-toluidin (BCIP) in $NaHCO_3$, pH 10.0.

**Live cell imaging**

Time-lapse imaging of cells was performed using an Olympus XI-81 inverted microscope (Olympus) controlled by Metamorph software (Molecular Devices) and equipped with objectives specified below and a CoolSnap EZ camera (Photometrics). Cells were seeded onto 35 mm glass bottom dishes (Ibidi) coated with 25 $\mu g \cdot ml^{-1}$ laminin (Sigma), and maintained in imaging medium composed of F-12 Ham Nutrient Mixture with 25 mM HEPES (Sigma) to compensate for the lack of $CO_2$, and supplemented with 10% FBS (Biowest), 1% Penicillin-Streptomycin (Biowest), 2 mM stable L-glutamine (Biowest), and 2.7 $g \cdot ml^{-1}$ L D-glucose (Carl Roth) in an Ibidi Heating System at 37°C. For random motility assays, cells were seeded at low density onto dishes and allowed to adhere for 3 h. Subsequently, the growth medium was exchanged with imaging medium, the chamber mounted into the heating system, and cells recorded by phase-contrast, time-lapse imaging at 1 min intervals for 3 h using an Uplan FL N 4x/0.13NA objective (Olympus) with 1.6x optovar intermediate magnification. Single cell tracking was performed in ImageJ by MTrackJ. Analyses of cell speed and cell trajectories, turning angles, and mean square displacements were performed in Excel (Microsoft) using a customized macro (4). Cells that touched each other or divided were excluded from analyses. Directionality index was determined by dividing the shortest distance between starting and end points (d) by the actual trajectories (D). Protrusion of lamellipodia was recorded at 5 s intervals for 10 min using an UPlan FI 100x/1.30NA oil immersion objective (Olympus). Protrusion rates of advancing lamellipodia assessed over a period of 2.5 min were quantified by generating kymographs from time-lapse movies of the cell periphery using Fiji, followed by slope determination from these kymographs. Lamellipodial persistence of randomly migrating cells on laminin was determined by phase-contrast, time-lapse microscopy using an UPlan FL N 100x/1.30NA objective and a frame rate of 1 frame per minute. Lamellipodial persistence was defined as time in min from initiation until collapse of the lamellipodium. For epifluorescence imaging of fluorescently labelled cells, B16-F1 and derived mutant cells were transfected with pEGFP-Lifeact. 16 h post transfection, the cells were seeded onto laminin-coated glass bottom dishes, and maintained 3 h in imaging medium. Migrating cells were recorded by time-lapse imaging at 10 s intervals for 10 min using an Olympus 100x/1.3NA Uplan FL objective. For multi-color TIRF imaging of fluorescently labelled cells, B16-F1 and derived mutant cells were co-transfected with pmCherry-MyoX and pEGFP-Lifeact. After seeding and adaptation in imaging medium for 3 h, the cells were recorded with a Nikon Eclipse TI-E inverted microscope equipped with a TIRF Apo 100x objective and Ixon3 897 EMCCD cameras (Andor) at 3 s intervals for 15 min.

**Immunofluorescence**

For immunofluorescence labelling, cells were fixed for 20 min in pre-warmed PBS, pH 7.3 containing 4% PFA and 0.06% picric acid, subsequently washed three times with PBS supplemented with 100 mM glycine. The cells were then permeabilized with 0.1% Triton X-100 in PBS for 30 s and blocked with PBG (PBS, 0.045% cold fish gelatin (Sigma), and 0.5% BSA). For immunolabeling with fascin, cells were fixed in -20 °C cold methanol. Primary antibodies were incubated overnight, followed by extensive washing of the specimens with PBG and incubation with respective secondary antibodies for at least 2 h. F-actin was visualized with Atto550 or Atto633-phalloidin. Imaging of fixed cells was conducted with an Olympus XI-81 inverted microscope equipped with an UPlan FI 100x/1.30NA oil immersion objective or a Zeiss LSM 980 equipped with Plan-Apochromat 63X/1.4 NA oil DIC objective using 488 nm, 594 nm and 633 nm laser lines. Fluorescence intensities of phalloidin or lamellipodial proteins were measured from still images captured at identical settings using Fiji/ImageJ software after background subtraction. Mean pixel intensities in lamellipodial regions of interest are shown as whiskers-box plots including all data points.

The Fiji plugin Coloc2 was used to calculate the co-localization of respective protein pairs in leading edge clusters. For this purpose, immunostained cells were recorded by confocal z-stack imaging followed by 3D reconstruction of data sets in Fiji using maximal intensity projections. Pearson correlation coefficients (r) were determined with the Coloc2 plugin to evaluate the co-localization of protein pairs in a 1 μm wide sector along leading edge. The co-localization of pixel intensities in two channels was visualized in cytofluorograms generated by a MATLAB exchange file (Timothy Olson, 2022; https://www.mathworks.com/matlabcentral/fileexchange/68712-cytofluorogram).

**Induction of filopodia in absence of lamellipodia**

B16-F1 wild-type and 4x-KO I-BAR cells were seeded onto low laminin (1 $\mu g \cdot ml^{-1}$) for 1 hr, a concentration lower than the threshold for inducing prominent lamellipodia in this cell type and conditions. Correspondingly, NIH 3T3 and derived 2xKO I-BAR cells were seeded onto low fibronectin (1 $\mu g \cdot ml^{-1}$) for 1 hr. Then, cells were incubated for 2 hr with the Arp2/3 complex inhibitor CK666 (200 μM) (Sigma) to abolish lamellipodia and trigger filopodia formation. Subsequently, cells were fixed and stained with phalloidin for the actin cytoskeleton followed by confocal imaging.

**TIRF assays**

For TIRF bead assays 50 µl of SNAP-capture magnetic beads with a diameter of 2 µm (NEB #S9145Z) were washed twice with coupling buffer (20 mM HEPES pH 7.4, 150 mM NaCl, 2 mM DTT, 2 mM β-mercaptoethanol, 10% BSA) and then coated with 10 µM of SNAP-GCN4-Lpd$^{850-1250}$, -Lpd$^{850-1250}$(44A), -Abi1$^{331-424}$-WT or -Abi1$^{331-424}$-mut, respectively, for 16 h on a wheel rotator at 4°C. Subsequently, the beads were washed three times with coupling buffer and the beads split into two fractions. One fraction was incubated with coupling buffer and served as negative control, while the other was incubated with 0.5 µM soluble His-VASPDdGAB in coupling buffer for 16 h at 4°C on a wheel rotator. Subsequently, beads were washed five times with coupling buffer and then re-suspended into storage buffer (20 mM HEPES pH 7.4, 150 mM NaCl, 2 mM DTT, 2mM β-mercaptoethanol, 10% BSA and 60% glycerol) for direct use in experiments or storage at -20°C. TIRF assays were performed in TIRF buffer (20 mM imidazole pH 7.4, 1 mM $MgCl_2$, 50 mM KCl, 0.5 mM ATP, 1 mM EGTA, 20 mM β-mercaptoethanol, 15 mM glucose, 20 $\mu g \cdot ml^{-1}$ catalase, 100 $\mu g \cdot ml^{-1}$ glucose oxidase, and 2.5 $mg \cdot ml^{-1}$ methylcellulose (4,000 cP)). Actin was polymerized in the presence of 25 nM CP and coated beads. Actin assembly was initiated by addition of G-actin (1 µM, 10% Atto488-labelled at Cys374) and the mixtures subsequently flushed into mPEG-silan (Mr 2,000) (Lysan Bio)-pre-coated flow chambers. Images were acquired with a Nikon Eclipse TI-E inverted microscope equipped with a TIRF Apo 100x objective at 3 s intervals with exposure times of 70 ms by Ixon3 897 EMCCD cameras (Andor) for at least 15 min.

**Adaptive kymographs and analyses of lateral MyoX cluster speed**

TIRF-microscopy, time-lapse movies of representative B16-F1 wild-type and EVM-KO mutant cells expressing EGFP-MyoX with a stable and constantly advancing lamellipodium were analyzed. A reference line in the direction of forward movement, but perpendicular to the lamellipodium was drawn. Guided by the reference line, line-scan measurements were performed using an 8 px wide and 100 px long line continuously tracking the advancing leading edge in consecutive frames using Fiji. For quantification of lateral MyoX cluster movement in the leading edge, stacks were re-oriented using the transformation function in Fiji to obtain horizontally positioned lamellipodia. After inserting a reference line, positioned parallel to the advancing cell front, the MTrackJ plug-in was used to track the lateral movement of clusters in consecutive frames, to determine average lateral cluster velocities. Data were processed and visualized using Excel and Origin software.

**BioID2 pulldown**

BioID2 assays were performed essentially as previously reported (6). For large-scale pulldowns, $5x10^7$ cells per construct were incubated with 50 µM biotin in growth medium overnight. After trypsinization, the cells were pelleted, washed twice with cold PBS and lysed in cold RIPA buffer supplemented with 0.4% sodium dodecyl sulfate (SDS), 5 mM benzamidine and 0.1 mM AEBSF for 30 min at 4°C on a wheel rotator. Subsequently,

Triton X-100 was added to a 2% final concentration followed by incubation for 30 min at 4°C on a wheel rotator. After addition of an equal volume of 50 mM Tris, pH 7.4, the lysate was cleared by centrifugation at 16,000xg and 4°C for 15 min. The supernatant was collected and incubated with 500 µl of Dynabeads (Invitrogen) on a rotary wheel overnight. Beads were then collected on a magnetic stand, and washed twice with wash buffer 1 (2% SDS in $H_2O$), once with wash buffer 2 (0.1% deoxycholate, 1% Triton X-100, 500 mM NaCl, 1 mM EDTA, and 50 mM HEPES, pH 7.5), once with wash buffer 3 (250 mM LiCl, 0.5% NP-40, 0.5% deoxycholate, 1 mM EDTA, and 10 mM Tris, pH 8.1) and finally twice with wash buffer 4 (50 mM Tris, pH 7.4, and 50 mM NaCl). 10% of the sample were used for analytic SDS-PAGE followed by silver staining. Bound proteins were removed from beads with SDS sample buffer supplemented with 50 µM biotin at 98°C. The remaining 90% of the sample were washed twice with 50 mM $NH_4HCO_3$ and used for mass spectrometry. Two independent large-scale BioID experiments with each construct were performed and isolated proteins from respective experiments pooled to increase protein yield for subsequent analysis.

**Protein Analyses by LC-MS**

Proteins from BioID pull downs were reduced with 5 mM DTT at 37°C for 1 h and alkylated with 10 mM iodoacetamide at RT for 30 min. Alkylation was quenched by adding DTT to a final concentration of 5 mM. Proteins bound to beads were then digested at 37 °C for 4 h using 1:100 w/w Lys-C (Wako) followed by overnight digestion with 1/100 w/w trypsin (Promega). Digestion was stopped by addition of trifluoroacetic acid (TFA) to a final concentration of 1%. Peptide solutions were then desalted with Sep Pak C18 1cc cartridges (Waters). Samples were dissolved in 0.1% TFA/2% acetonitrile and analyzed in an Orbitrap Fusion Lumos mass spectrometer (Thermo Fisher Scientific) equipped with a nanoelectrospray source and connected to an Ultimate 3000 RSLC nanoflow system (Thermo Fisher Scientific) with standard adjustments for peptide analysis in data-dependent acquisition mode (13). Raw MS data were processed using Max Quant software version 1.5 (14) , Perseus software version 1.6.2.3 (15) and mouse entries of Uniprot data base. Proteins were stated identified using a false discovery rate of 0.01 on protein and peptide level.

**Statistical analyses**

Quantitative experiments were performed at least in triplicates to avoid any potential environmental bias or unintentional error. Lamellipodial phenotypes as derived from analyses on fixed samples or living cells were systematically obtained from sample sizes of dozens or hundreds of cells, respectively. Raw data were processed in Excel (Microsoft). Statistical analyses were performed with Origin 2021 (OriginLab). All data sets were tested for normality by the Shapiro-Wilk test. Statistical differences between normally distributed datasets of two groups were determined by t-test and not normally distributed datasets of two groups by non-parametric Mann-Whitney U rank sum test. For comparison of more than two groups, statistical significance of normally distributed data was examined by one-way ANOVA and Tukey Multiple Comparison test. In case of not normally distributed data, the non-parametric Kruskal-Wallis test and Dunn’s Multiple Comparison test were used. Statistical differences were defined as *$p \leq 0.05$, **$p \leq 0.01$, ***$p \leq 0.001$ as well as n.s., not significant, and are displayed and stated in figures and figure legends, respectively.

### Supplementary Figures

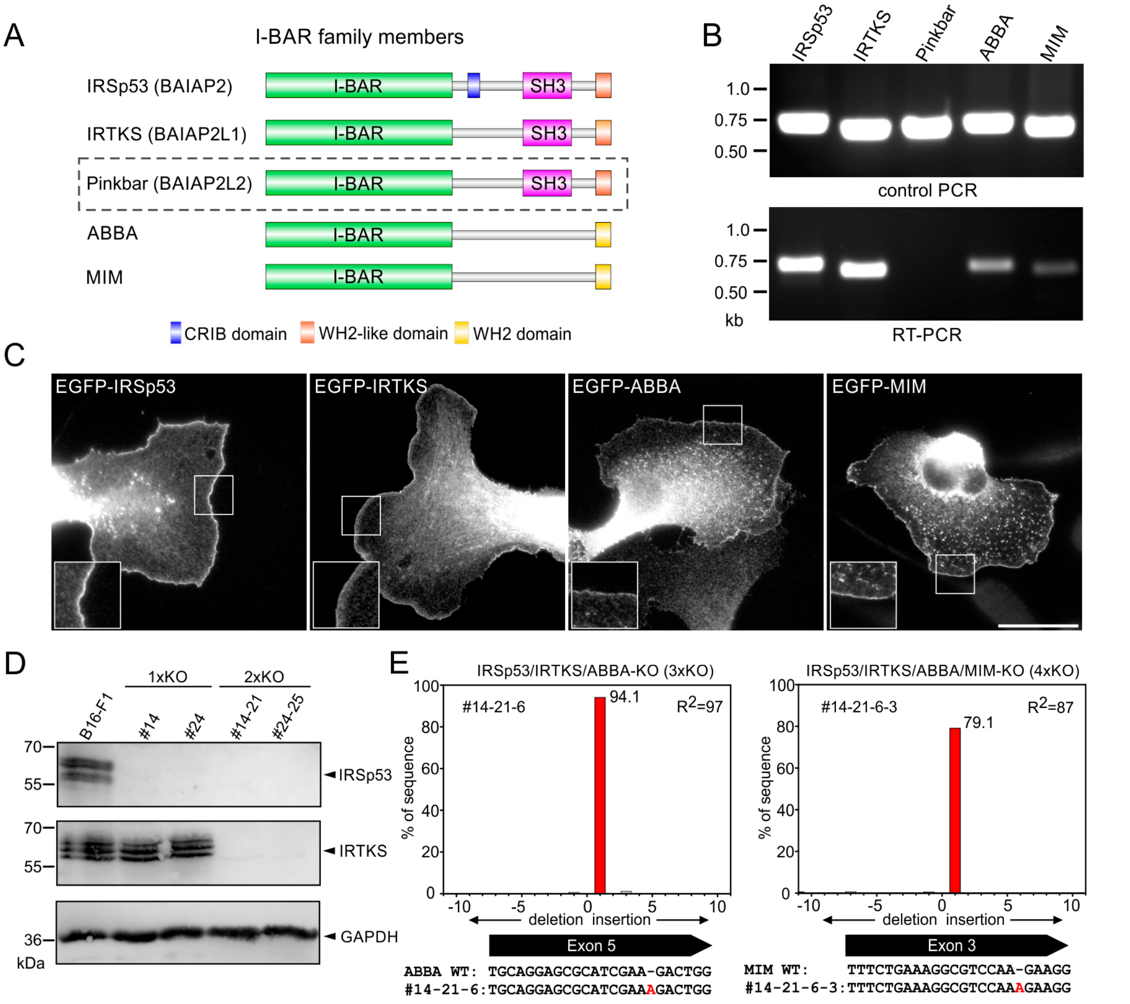


**Fig. S1. Generation of B16-F1-derived mutant cell lines lacking I-BAR protein expression.** (*A*) Domain organization of the five known I-BAR family members IRSp53 (BAIAP2), IRTKS (BAIAP2L1), Pinkbar (BAIAP2L2), ABBA (MTSS2) and MIM (MTSS1). In contrast to IRSp53, IRTKS and Pinkbar, both ABBA and MIM lack SRC Homology 3 (SH3) domains. IRSp53 additionally contains an unconventional Cdc42- and Rac-interactive binding (CRIB) motif. (*B*) Control PCRs using plasmid templates harboring respective full-length cDNAs encoding all I-BAR family proteins are shown on top panel. Semi quantitative RT-PCR using the same primer pairs and B16-F1 cDNA as template are shown at the bottom. (*C*) Ectopically expressed IRSp53, IRTKS, ABBA and

MIM fused to EGFP localize to the leading edge and intracellular vesicles in B16-F1 cells migrating on laminin. Insets, enlarged images of boxed regions. Bar, 15 µm. (*D*) Immunoblot confirming the elimination of IRSp53 in two independent single-knockout B16-F1 mutants (1xKO #14 and #24) and IRTKS in independent IRSp53/IRTKS double-knockout mutants (2xKO #14-21 and #24-25). Loading control: GAPDH. (*E*) Derived 3xKO (IRSp53/IRTKS/ABBA) mutant #14-21-6 or 4xKO (IRSp53/IRTKS/ABBA/MIM) mutant #14-21-6-3 additionally lacking ABBA and MIM, respectively, were identified by the TIDE sequence trace decomposition web tool (*top*) and confirmed by sequencing of respective target sites (*bottom*). $R^2$ indicates goodness of fit. Shown below are the homozygous frame-shift mutations in the 3xKO and 4xKO clones that were mainly used in this study.

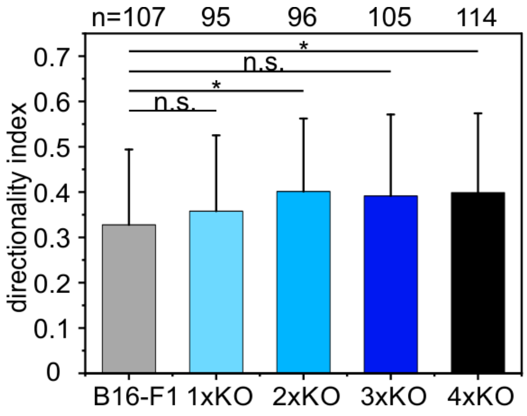


**Fig. S2. Loss of I-BAR proteins increases directionality of migrating cells.** Directionality of randomly migrating cells on laminin increased with consecutive inactivation of IRSp53 and IRTKs and then remained constant. Bars represent arithmetic means ± SD. Non-parametric Kruskal-Wallis test was used to reveal statistically significant differences between datasets. *p≤0.05; n.s.; not significant. n, number of cells. Pooled data from at least three independent experiments are shown.

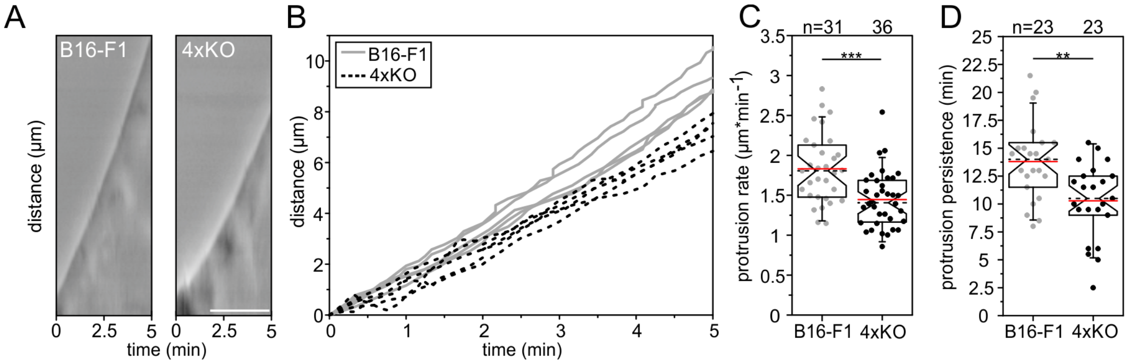


**Fig. S3.** (*A*) Loss of I-BAR proteins reduces the efficacy of lamellipodium protrusion. Kymographs of representative phase-contrast movies are shown. Bar, 2.5 µm. (*B*) Multiple examples of lamellipodium protrusion in B16-F1 *versus* 4xKO I-BAR cells (clone #14-21-6-3). (*C*) Quantification of protrusion rates. (*D*) Quantification of protrusion persistence. (C-D) Notched boxes in box plots indicate 50% (25-75%) and whiskers 90% (5-95%) of all measurements. Arithmetic means are highlighted in red and dashed black lines depict medians. (C-D) parametric, t-test were used to reveal statistically significant differences between datasets. **p≤0.01; ***p≤0.001; n, number of cells. C-D show pooled data from at least three independent experiments.

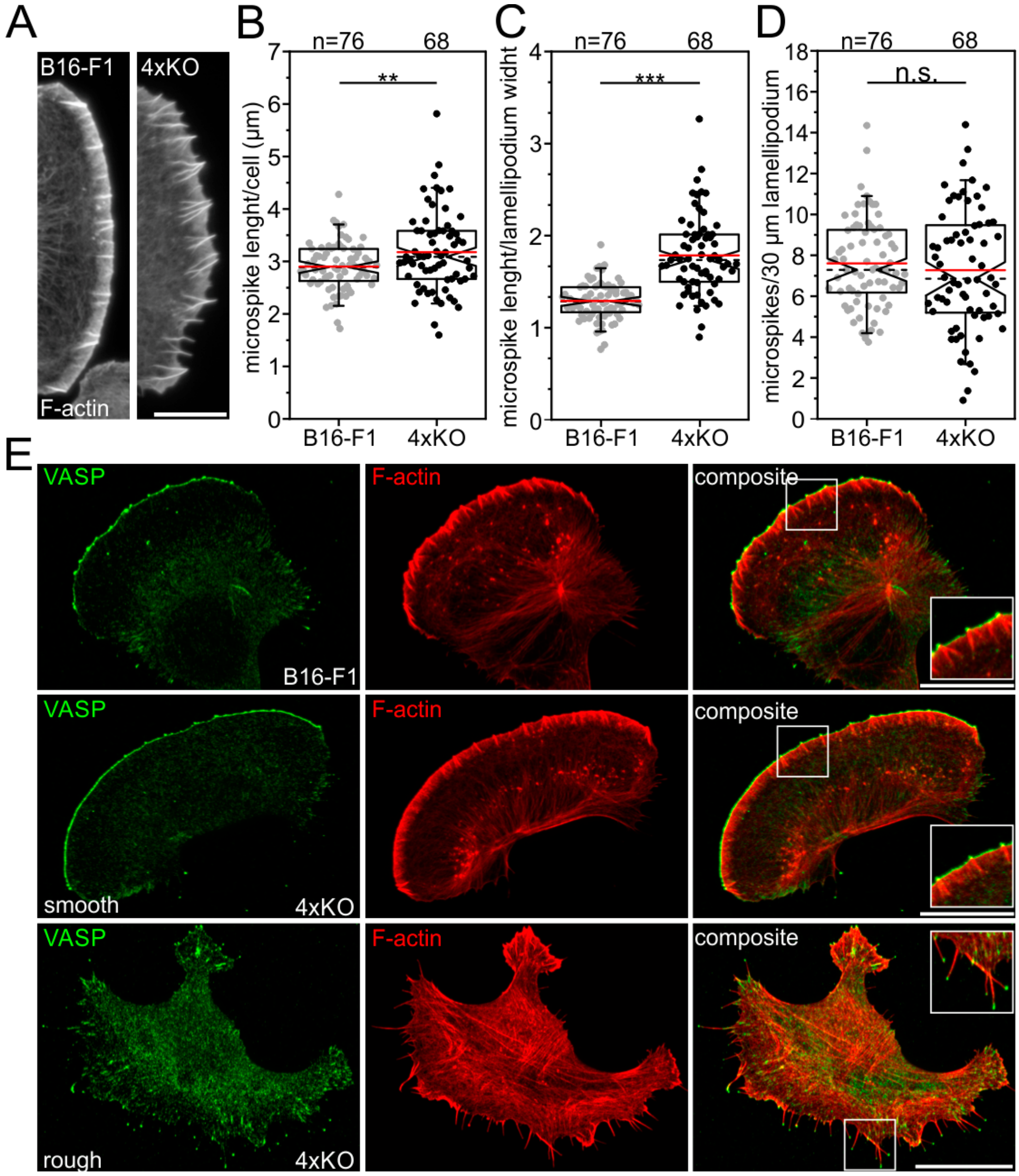


**Fig. S4. Loss of all I-BAR proteins does not impair formation of microspikes and filopodia.** (*A*) Representative images of microspikes in B16-F1 and 4xKO I-BAR cells. Bar, 10 µm. (*B*) Quantification of microspike length per cell. (*C*) Quantification of microspike length per cell over lamellipodium width. (*D*) Quantification of microspike numbers. (*E*) Despite loss of all expressed I-BAR family proteins, the 4xKO mutant still formed VASP clusters at the tips of microspikes (*middle*) and filopodia (*bottom*). Galleries display B16-F1 and 4xKO I-BAR cells stained for F-actin and endogenous VASP. Bars, 20 µm. Insets, enlarged images of boxed regions. (B-D) Notched boxes in box plots indicate 50% (25-75%) and whiskers 90% (5-95%) of all measurements; arithmetic means are highlighted in red and dashed black lines are depicting the medians. (B-C) t-test or (D) Mann-Whitney-U-test were used for statistical analysis. ***$p \leq 0.001$; n.s.; not significant. n, number of cells. B-D show pooled data from three independent experiments.

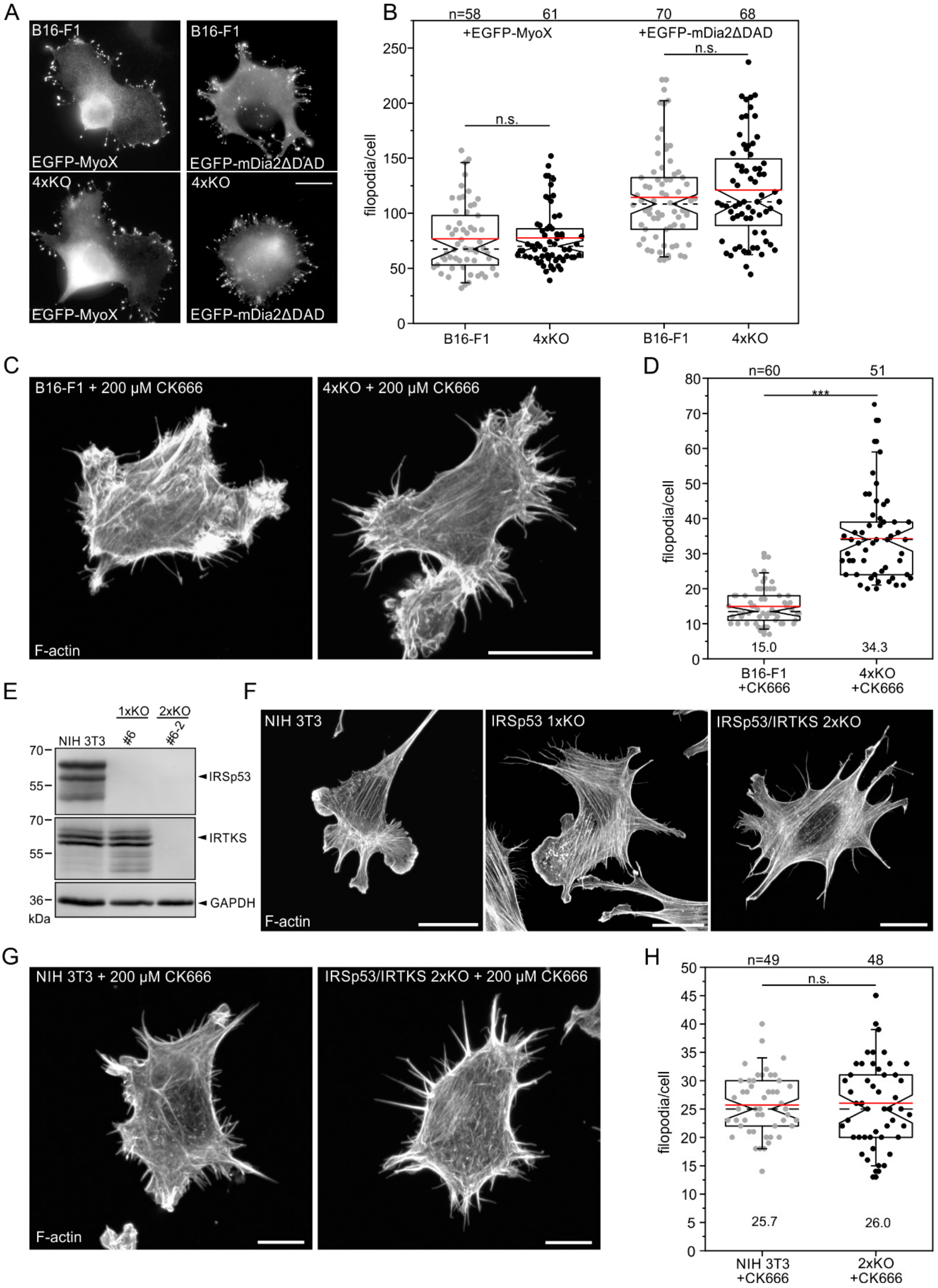
A
B16-F1
EGFP-MyoX
B16-F1
EGFP-mDia2ΔDAD
4xKO
EGFP-MyoX
4xKO
EGFP-mDia2ΔDAD
B
n=58
61
70
68
+EGFP-MyoX
+EGFP-mDia2ΔDAD
n.s.
n.s.
filopodia/cell
B16-F1
4xKO
B16-F1
4xKO
C
B16-F1 + 200 µM CK666
4xKO + 200 µM CK666
F-actin
D
n=60
51
***
filopodia/cell
15.0
34.3
B16-F1 +CK666
4xKO +CK666
E
1xKO
2xKO
NIH 3T3
#6
#6-2
IRSp53
IRTKS
GAPDH
kDa
F
NIH 3T3
IRSp53 1xKO
IRSp53/IRTKS 2xKO
F-actin
G
NIH 3T3 + 200 µM CK666
IRSp53/IRTKS 2xKO + 200 µM CK666
F-actin
H
n=49
48
n.s.
filopodia/cell
25.7
26.0
NIH 3T3 +CK666
2xKO +CK666

**Fig. S5. Loss of all I-BAR proteins does not impair formation of filopodia in different cell types.** (*A*) Unchanged filopodium formation in B16-F1 and 4x-KO (#14-21-6-3) cells induced by transient expression of myosin-X (myoX) or active mDia2 (mDia2ΔDAD), as indicated. Representative examples of B16-F1 and 4xKO cells after labelling with GFP nanobodies are shown. Bar, 20 µm. (*B*) Quantification of filopodia in B16-F1 and 4xKO mutant cells ectopically expressing MyoX or active mDia2. (*C*) 4xKO I-BAR cells form excess filopodia in the absence of lamellipodia. Representative examples of phalloidin-stained wild-type B16-F1 and 4xKO cells devoid of lamellipodia after treatment with 200 mM CK666 seeded at low laminin (1 µg $ml^{-1}$). Bar, 20 µm. (*D*) Quantification of filopodia in CK666-treated B16-F1 and 4xKO cells. (*E*) Generation of NIH 3T3-derived mutant cell lines lacking the two most abundant I-BAR family members IRSp53 and IRTKS. Immunoblot confirming the elimination of IRSp53 in the NIH 3T3 1xKO mutant (#6) and IRTKS in the IRSp53/IRTKS 2xKO mutant (#6-2). Loading control: GAPDH. (*F*) Representative images of wild-type and mutant cells as indicated and stained for F-actin. Bar, 20 µm. (*G*) 2xKO I-BAR cells form comparable numbers of filopodia in the absence of lamellipodia. Representative examples of phalloidin-stained wild-type NIH 3T3 and 2xKO mutant cells devoid of lamellipodia after treatment with 200 mM CK666 seeded at low fibronectin (1 µg $ml^{-1}$). Bar, 10 µm. (*H*) Quantification of filopodia in CK666-treated NIH 3T3 and 2xKO cells. (B, D and H) Notched boxes in box plots indicate 50% (25-75%) and whiskers 90% (5-95%) of all measurements; arithmetic means are highlighted in red and dashed black lines are depicting the medians. (B, D) Non-parametric, Mann-Whitney-U-test and (H) parametric, t-test were used to reveal statistically significant differences between datasets. ***p≤0.001; n.s.; not significant. n, number of cells. B, D and H show pooled data from three independent experiments.

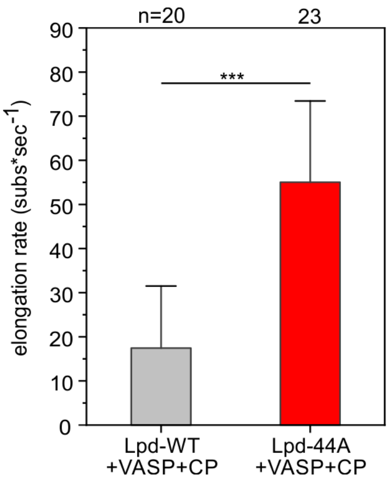


**Fig. S6. VASP-mediated actin filament elongation on Lpd-44A-derivatized beads is about 3-fold faster as compared to filament assembly on beads coated with Lpd-WT.** Quantification of filament elongation rates of processively growing filaments on SNAP-GCN4-Lpd$^{850-1250}$-derivatized beads in the presence of 25 nM CP after incubation with soluble VASP is shown. Data were analyzed under conditions and from movies as shown in Fig. 4b. Bars show means±SD of pooled data from three independent experiments. Non-parametric, Mann-Whitney-U-test was used to reveal statistically significant differences between datasets. ***p≤0.001. n, number of filaments analyzed.

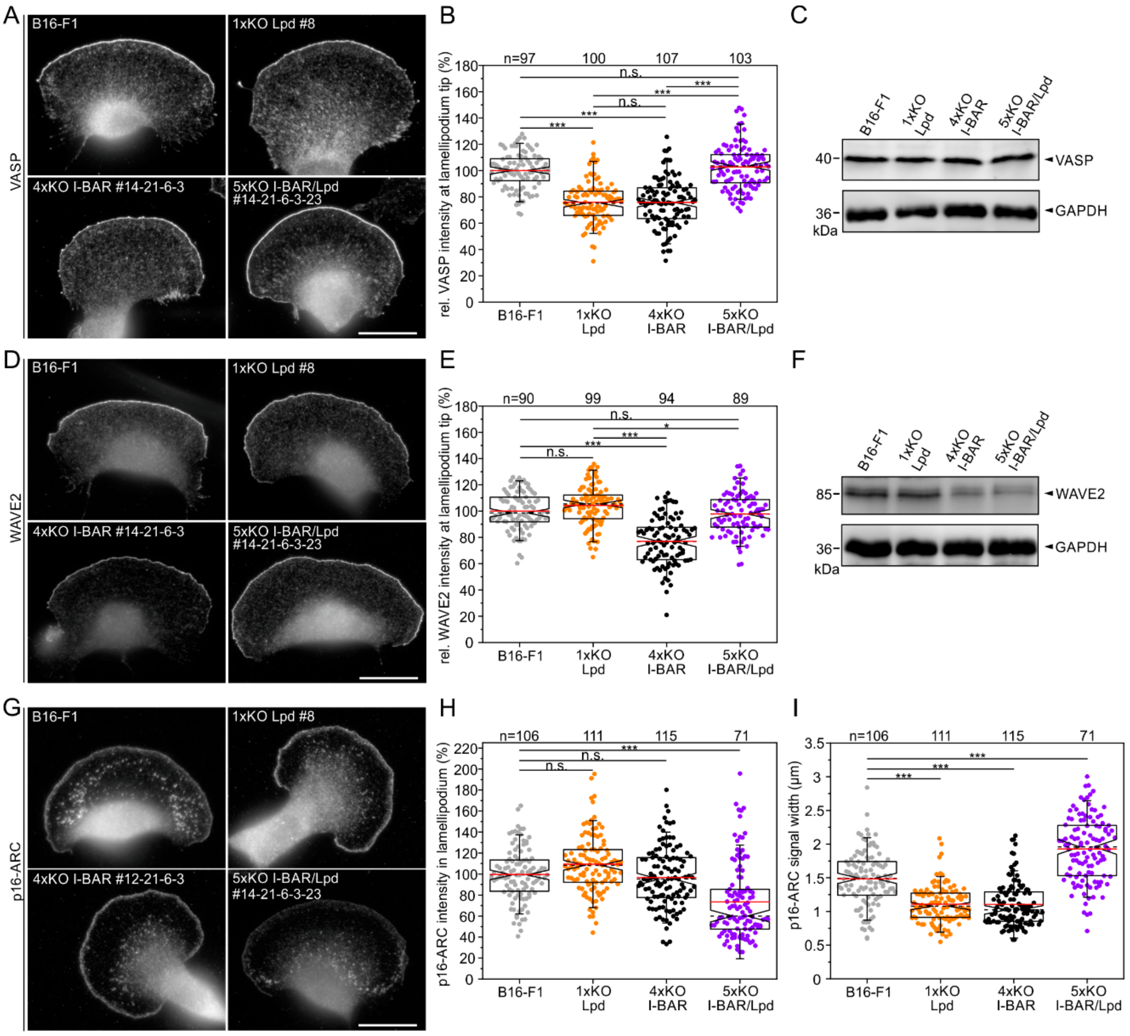


**Fig. S7. The combined loss of I-BAR proteins and Lpd compensates for the defects of the respective other regarding distribution of relevant lamellipodial marker proteins.** (*A*) Representative images of wild-type and mutant cells stained for VASP. Bar, 15 µm. (*B*) Quantifications of VASP intensities at lamellipodia tips from images as shown in A. (*C*) Comparable expression of VASP in wild-type and mutant cells was confirmed by immunoblotting. GAPDH was used as loading control. (*D*) Representative images of wild-type and mutants cells stained for WAVE2. Bar, 15 µm. (*E*) Quantifications of WAVE2 intensities at lamellipodia tips from images as shown in D. (*F*) Diminished expression of WAVE2 in mutant cells lacking I-BAR proteins was confirmed by immunoblotting.

GAPDH was used as loading control. (*G*) Representative images of wild-type and mutant cells stained for the Arp2/3 complex subunit p16-ARC. Bar, 15 µm. (*H*) Quantifications of p16-ARC intensities in lamellipodia from images as shown in G. (*I*) Quantification of p16-ARC signal width in lamellipodia. (B-D, F-G) Notched boxes in box plots indicate 50% (25-75%) and whiskers 90% (5-95%) of all measurements; arithmetic means are highlighted in red, with dashed black lines depicting the medians. (B, E) parametric, 1-way ANOVA test or (H, I) non-parametric, Kruskal-Wallis test were used to reveal statistically significant differences between datasets. ***$p \leq 0.001$; *$p \leq 0.05$; n.s.; not significant. n, number of cells. B, E, H, and I show pooled data from at least three independent experiments.

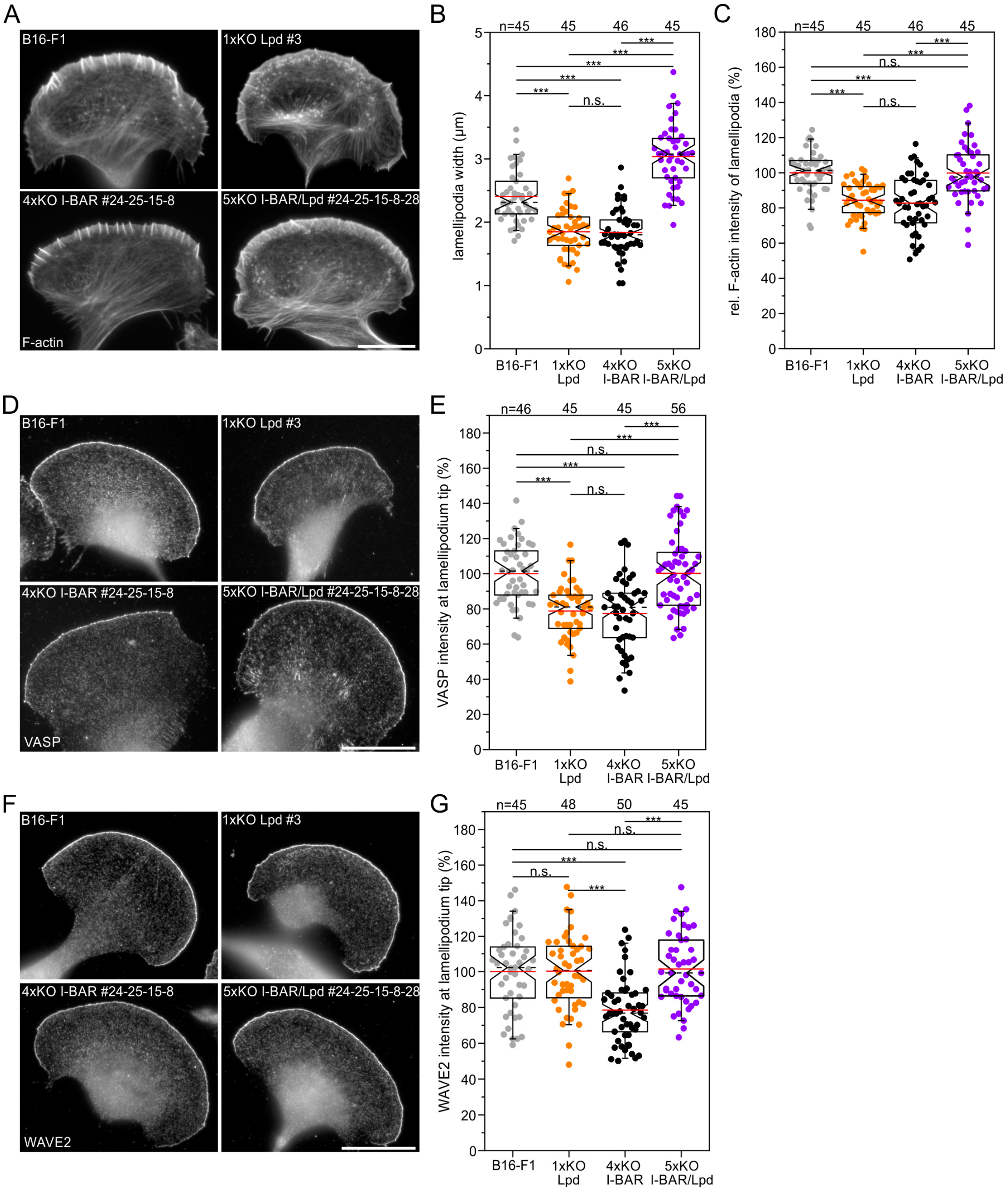


**Fig. S8. Validation of key lamellipodial parameters in independent set of clones lacking Lpd and I-BAR proteins.** (*A*) Representative images of wild-type and an independent set of mutant cells as indicated and stained for F-actin. Bar, 20 µm. (*B*) Quantification of lamellipodium width. (*C*) Quantification of F-actin intensity. (*D*)

Representative images of wild-type and mutants cells stained for VASP. Bar, 20 µm. (*E*) Quantifications of VASP intensities at lamellipodia tips from images as shown in D. (*F*) Representative images of wild-type and mutants cells stained for WAVE2. Bar, 20 µm. (*G*) Quantifications of WAVE2 intensities in lamellipodia from images as shown in F. Once again, the combined loss of I-BAR proteins and Lpd compensates for the defects of the respective other regarding distribution of relevant lamellipodial marker VASP and WAVE2. (B, C, E and G) Notched boxes in box plots indicate 50% (25-75%) and whiskers 90% (5-95%) of all measurements; arithmetic means are highlighted in red, with dashed black lines depicting the medians. Parametric, 1-way ANOVA test was used to reveal statistically significant differences between datasets. ***p≤0.001; n.s.; not significant. n, number of cells. B, C, E, and G show pooled data from three independent experiments.

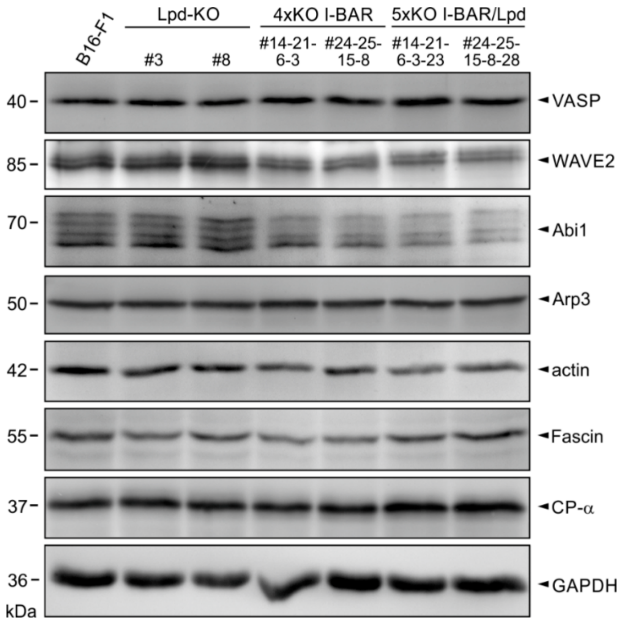


**Fig. S9. Global expression of relevant marker proteins in independent clonal cell lines.** Immunoblots of relevant lamellipodial marker proteins as indicated in two complete sets of independent mutants devoid of Lpd, I-BAR or a combination of both are shown. GAPDH was used as loading control. As opposed to expression of the other proteins that remained virtually unchanged, note the noticeably diminished, but mutually comparable, expression of the WRC components WAVE2 and Abi1 in all mutants lacking the I-BAR proteins.

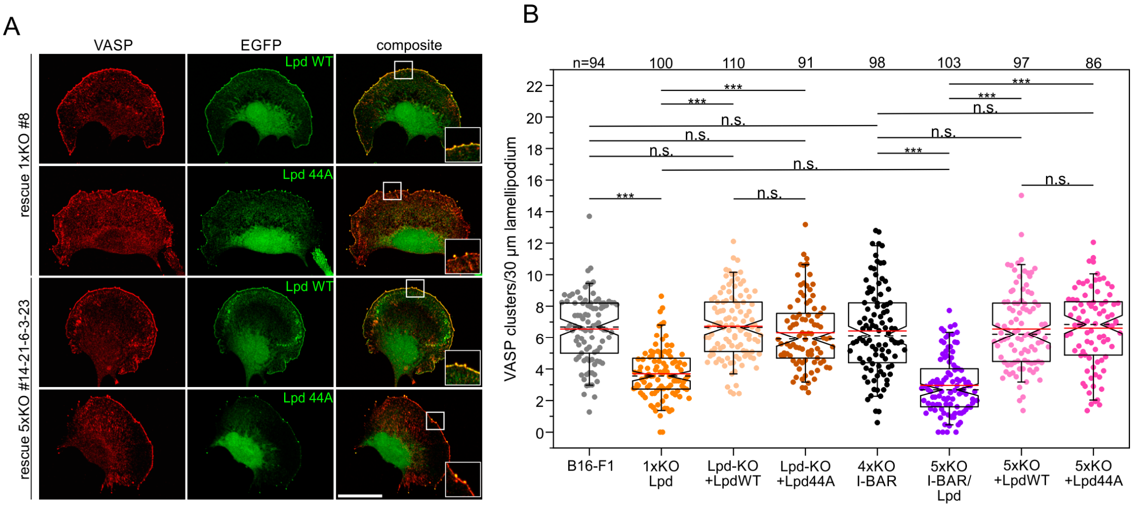


**Fig. S10. Lpd contributes to VASP clustering.** (*A*) Representative images of 1xKO Lpd and 5xKO I-BAR/Lpd mutant cells reconstituted with full-length Lpd-WT *versus* Lpd-44A fused to EGFP. Fixed cells were stained for endogenous VASP and EGFP signals enhanced with nanobodies. Insets, enlarged images of boxed regions. Bar, 20 µm. (*B*) Quantification of VASP clustering in wild-type, mutant and reconstituted mutant cells. Notched boxes in box plots indicate 50% (25-75%) and whiskers 90% (5-95%) of all measurements; arithmetic means are highlighted in red and dashed black lines depicting the medians. Kruskal-Wallis test was used for statistical analysis. ***$p \leq 0.001$; n.s.; not significant. n, number of cells. Pooled data from three independent experiments are shown.

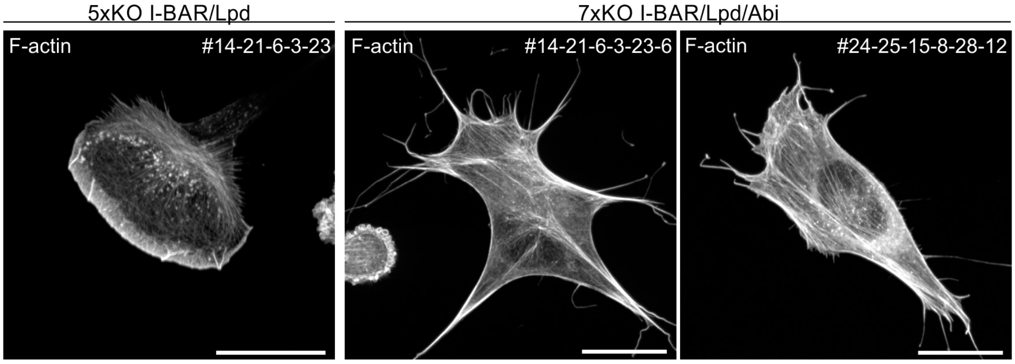


**Fig. S11. Combined inactivation of Abi1 and Abi2 in 5x-KO I-BAR/Lpd mutant cells abolishes lamellipodium formation.** Representative 5xKO and two independent 7xKO cells stained for the F-actin cytoskeleton with phalloidin are shown. Bars, 20 µm.

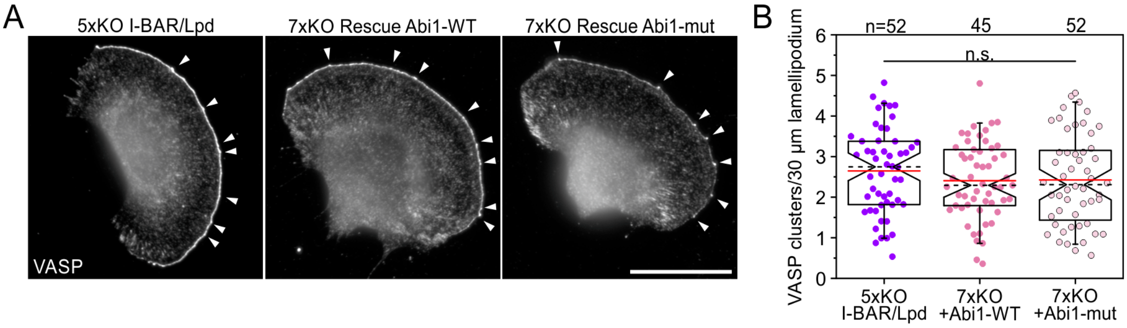


**Fig. S12. Loss of the VASP-Abi interaction does not significantly affect the clustering of VASP.** (*A*) Representative images of fixed 5xKO I-BAR/Lpd control and reconstituted 7xKO I-BAR/Lpd/Abi cells expressing full-length Abi1-WT *versus* Abi1-mut fused to EGFP and stained for endogenous VASP. White arrowheads indicate VASP clusters. Bar, 20 µm. (*B*) Quantification of VASP clustering in 5xKO control and reconstituted mutant cells as indicated. Notched boxes in box plots indicate 50% (25-75%) and whiskers 90% (5-95%) of all measurements; arithmetic means are highlighted in red and dashed black lines depicting the medians. Parametric, 1-way ANOVA test was used to reveal statistically significant differences between. n.s.; not significant. n, number of cells. Pooled data from three independent experiments are shown.

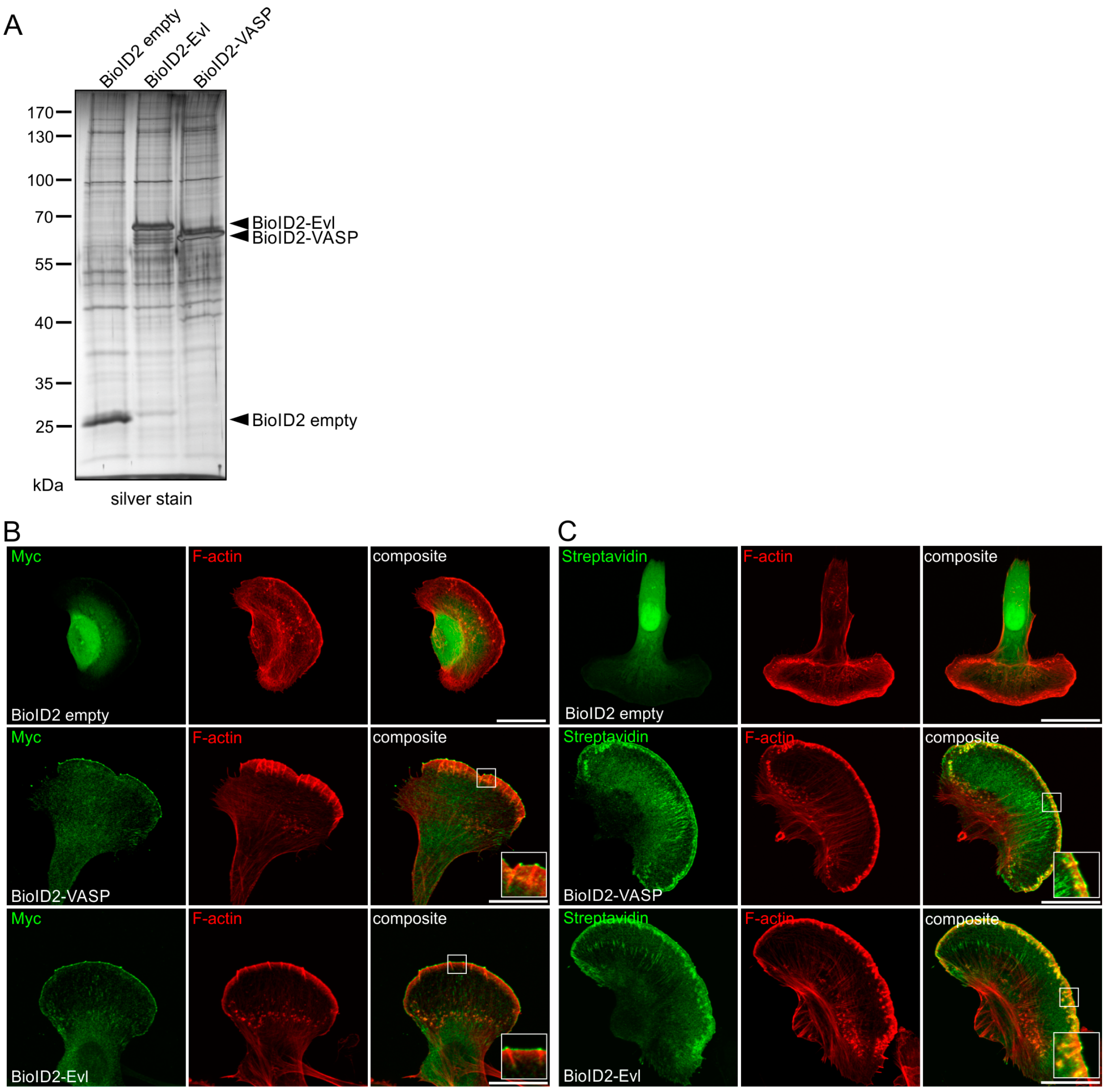


**Fig. S13. Validation of used BioID constructs.** (*A*) Empty BioID2-control as well as the BioID2-fusion protein baits indicated were expressed at comparable levels in Ena/VASP deficient EVM-KO cells. Analytical samples from BioID2 pulldowns on streptavidin-coated Dynabead pulldowns were separated by SDS-PAGE and subjected to silver staining. (*B*) Representative images of transfected EVM-KO cells expressing the constructs indicated and stained for Myc-tagged BioID constructs and F-actin. In contrast to the BioID-empty control that mainly accumulated in the nucleus of EVM-KO cells, BioID-tagged VASP and Evl were recruited to the tips of the leading edge and rescued microspikes as well as to focal adhesions. (*C*) Comparable images of transfected EVM-KO cells stained for biotinylated proteins with fluorescent streptavidin and for F-actin. (B and C) Bars, 20 µm. Insets, enlarged images of boxed regions.

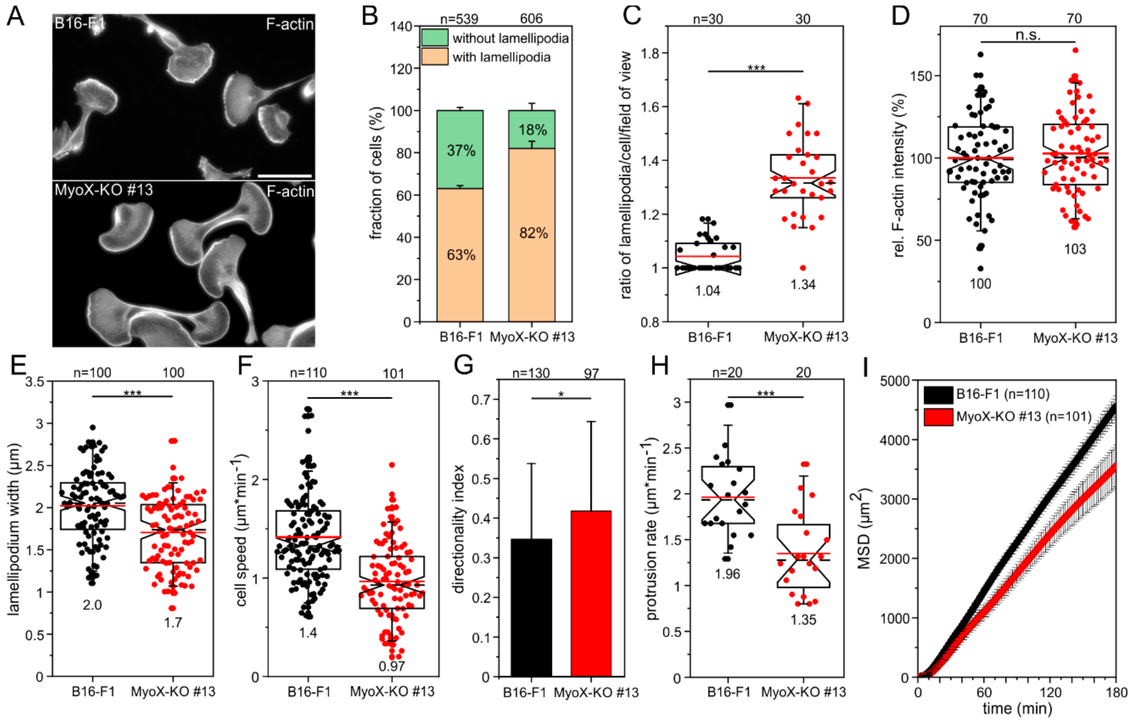


**Fig. S14. Loss of MyoX impairs cell migration.** (*A*) Formation of multiple fronts in MyoX-KO cells as visualized with phalloidin. Bar, 50 µm. (*B*) Quantification of lamellipodia. Bars represent arithmetic means ± SD from at least three independent experiments; n, number of analyzed cells. (*C*) Quantification of protrusive fronts. Error bars represent means ± SEM from at least three independent experiments. (*D-G*) Quantifications of F-actin intensities (*D*), lamellipodium width (*E*), 2D migration on laminin (*F*) and directionality (*G*). Bars in the latter represent means ± SD. (*H*) Quantification of protrusion rates. (*I*) Analyses of MSD. Respective symbols and error bars represent means ± SEM. Notched boxes in box plots in C-F and H indicate 50% (25-75%) and whiskers 90% (5-95%) of all measurements; arithmetic means are highlighted in red and dashed black lines depicting the medians. (D and H) Parametric, t-test, (C, E-G) or non-parametric, Mann-Whitney-U-test were used to reveal statistically significant differences between data sets. *p≤0.05; ***p≤0.001; n.s.; not significant. n, number of analyzed cells (B, D-I) or number of analyzed images with more than 340 cells per cell line (C). B-I show pooled data from at least three independent experiments.

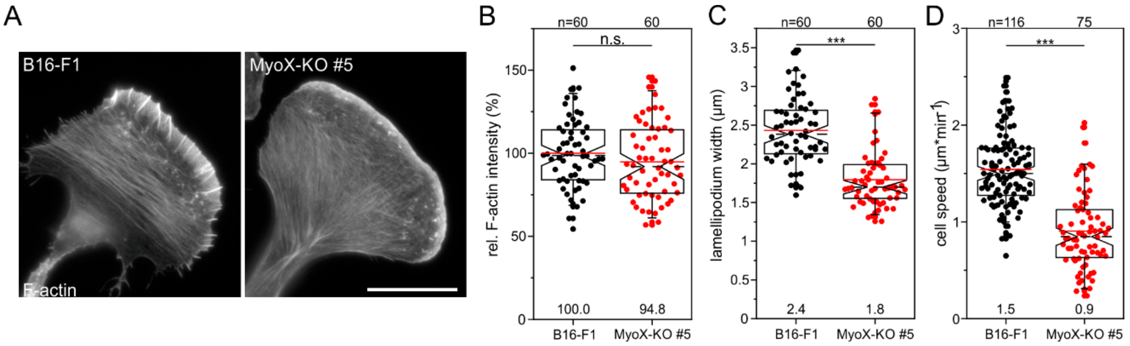


**Fig. S15. Validation of lamellipodium characteristics and 2D-random cell migration in an independent clone lacking MyoX.** (*A*) Representative images of a B16-F1 wild-type and derived MyoX-KO mutant (clone #5) cell stained for F-actin. Comparably to MyoX-clone #13, this clone also completely lacked microspikes. Bar, 20 µm. (*B*) Quantification of F-actin intensity. (*C*) Quantification of lamellipodium width. (*D*) Quantification of 2D migration on laminin. Notched boxes in box plots in B-D indicate 50% (25-75%) and whiskers 90% (5-95%) of all measurements; arithmetic means are highlighted in red and dashed black lines depicting the medians. (B) Parametric, t-test or

(C and D) non-parametric, Mann-Whitney-U-test were used to reveal statistically significant differences between data sets. ***p≤0.001; n.s.; not significant. n, number of analyzed cells. B-D show pooled data from at least three independent experiments.

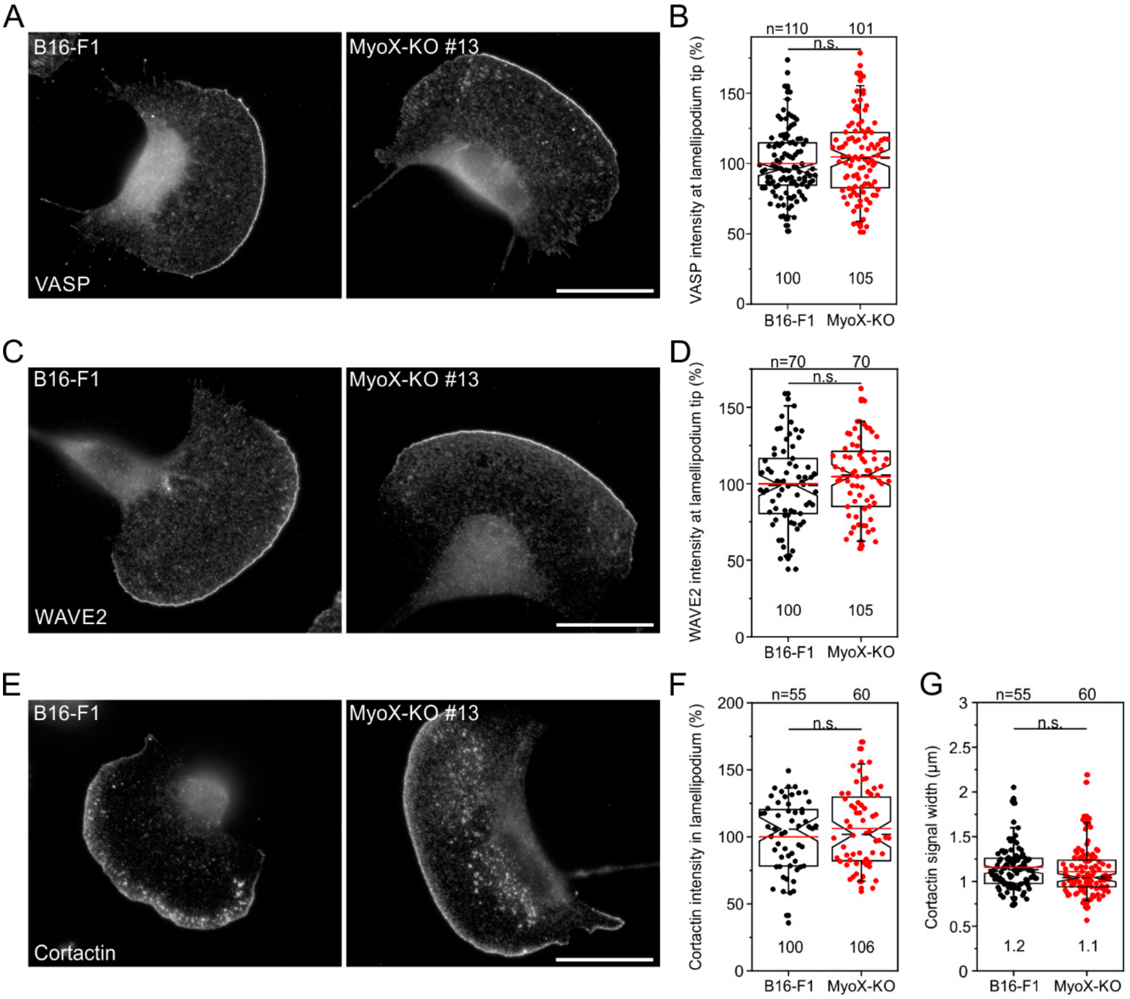


**Fig. S16. Loss of MyoX does not affect distribution of prominent lamellipodial marker proteins.** (*A, C, E*) Representative images of B16-F1 wild-type and MyoX-KO cells stained for VASP, WAVE2 and cortactin, as indicated. Bars, 20 µm. (*B, D, F*) Quantifications of respective protein intensities in lamellipodia from images as shown in A. (*G*) Quantification of cortactin signal width in lamellipodia. (B-D, F-G) Notched boxes in box plots indicate 50% (25-75%) and whiskers 90% (5-95%) of all measurements, arithmetic means are highlighted in red, with dashed black lines depicting the medians. (D, F) Parametric, t-test or (B, G) Non-parametric, Mann-Whitney-U-test were used to reveal statistically significant differences between datasets. n.s.; not significant. n, number of cells. B, D, and F-G show pooled data from three independent experiments.

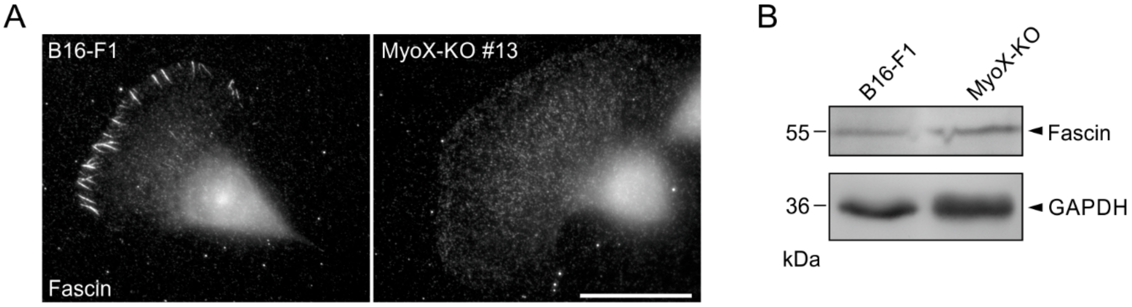


**Fig. S17.** (*A*) Loss of MyoX abolishes fascin-decorated microspikes. Immunostaining of methanol-fixed, wild-type and MyoX-deficient cells with fascin antibody is shown. Bar, 20 µm. (*B*) Comparable expression of fascin in B16-F1 wild-type and MyoX-KO cells was confirmed by immunoblotting using fascin-specific antibodies. GAPDH was used as loading control.

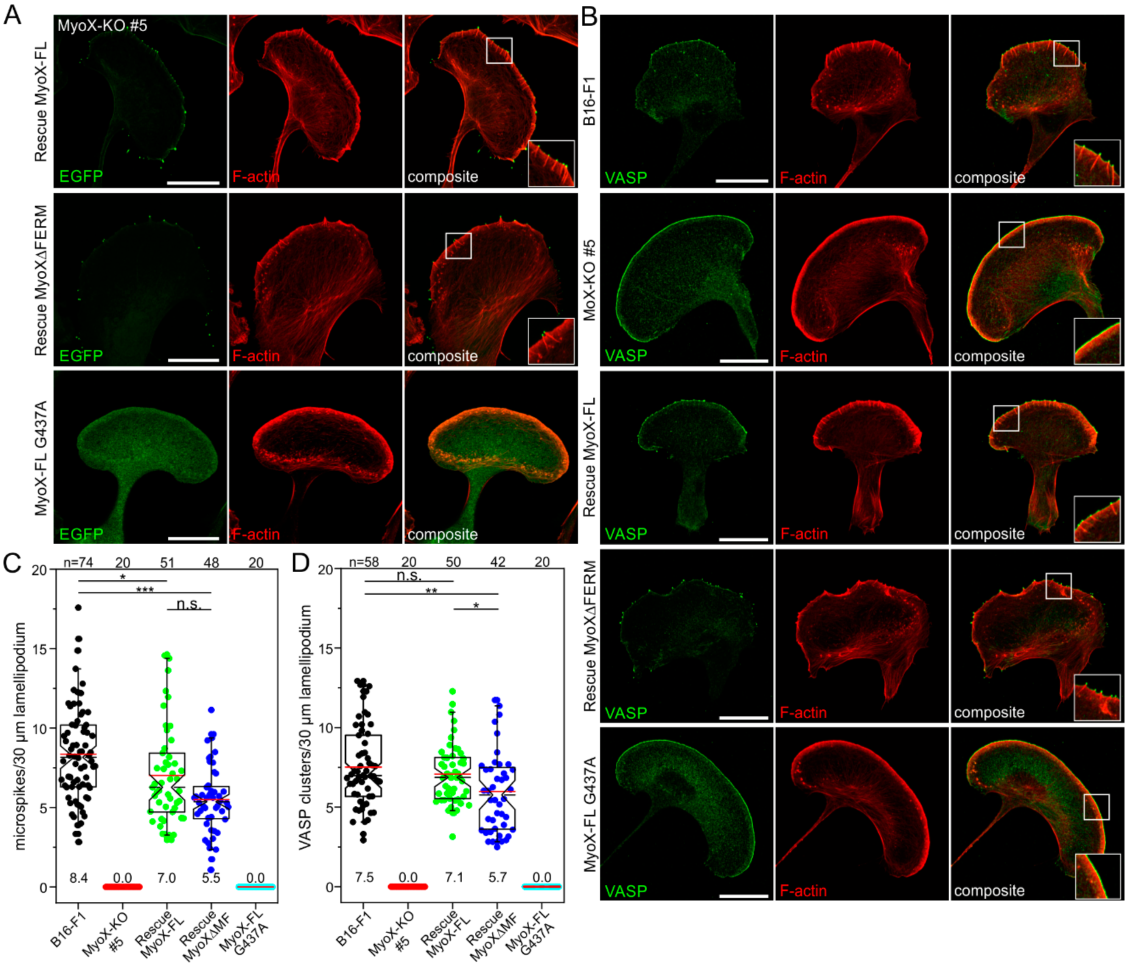


**Fig. S18. Loss of MyoX eliminates microspike formation due to abolished VASP clustering in independent MyoX-KO clone #5.** (*A*,*B*) Ectopically expressed full-length MyoX and MyoXΔMF, but not the MyoX motor dead variant (G437A), in the MyoX-KO mutant rescue microspike formation (A) and VASP clustering (B). Fixed cells were stained for F-actin and EGFP signals enhanced with nanobodies (A), or for F-actin and endogenous VASP (B). Insets, enlarged images of boxed regions. Bars, 20 µm. (*C*) Quantification of microspikes in wild-type, mutant and reconstituted cells. (*D*) Quantification of VASP clustering in wild-type, mutant and reconstituted cells. Notched boxes in box plots indicate 50% (25-75%) and whiskers 90% (5-95%) of all measurements; arithmetic means are highlighted in red and dashed black lines depicting the medians. (C, D) Statistical analysis by Kruskal-Wallis test. *p≤0.05; **p≤0.01; ***p≤0.001; n.s.; not significant. n, number of analyzed cells. C and D show pooled data from three independent experiments.

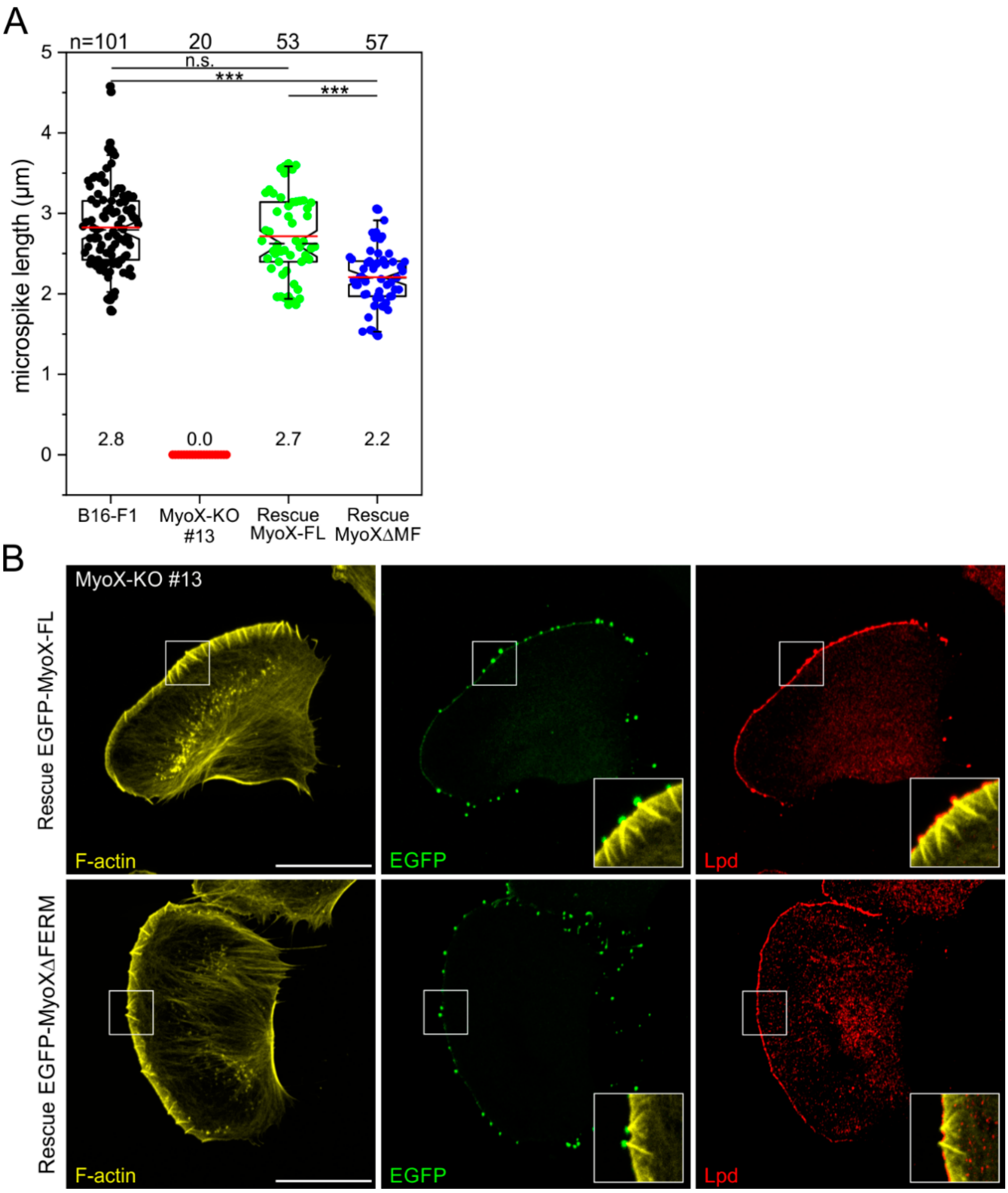


**Fig. S19. Reconstituted MyoX-KO cells expressing MyoXΔMF form shorter microspikes that fail to recruit Lpd to their tips.** (*A*) Quantification of microspike length in wild-type, mutant and reconstituted cells expressing full-length MyoX *versus* MyoXΔMF. Notched boxes in box plots indicate 50% (25-75%) and whiskers 90% (5-95%) of all measurements; arithmetic means are highlighted in red and dashed black lines depict the medians. Non-parametric Kruskal-Wallis test was used to reveal statistically significant differences between datasets. ***$p \leq 0.001$; n.s.; not significant. n, number of analyzed cells. Pooled data from three independent experiments are shown. (*B*) Comparison of reconstituted MyoX-KO cells expressing either full-length MyoX or MyoXΔMF fused to EGFP. Fixed cells were stained for F-actin, endogenous Lpd and EGFP signals enhanced with nanobodies. Note complete absence of Lpd in microspike tip clusters containing MyoXΔMF. Insets, enlarged composite images of boxed regions. Bars, 20 μm.

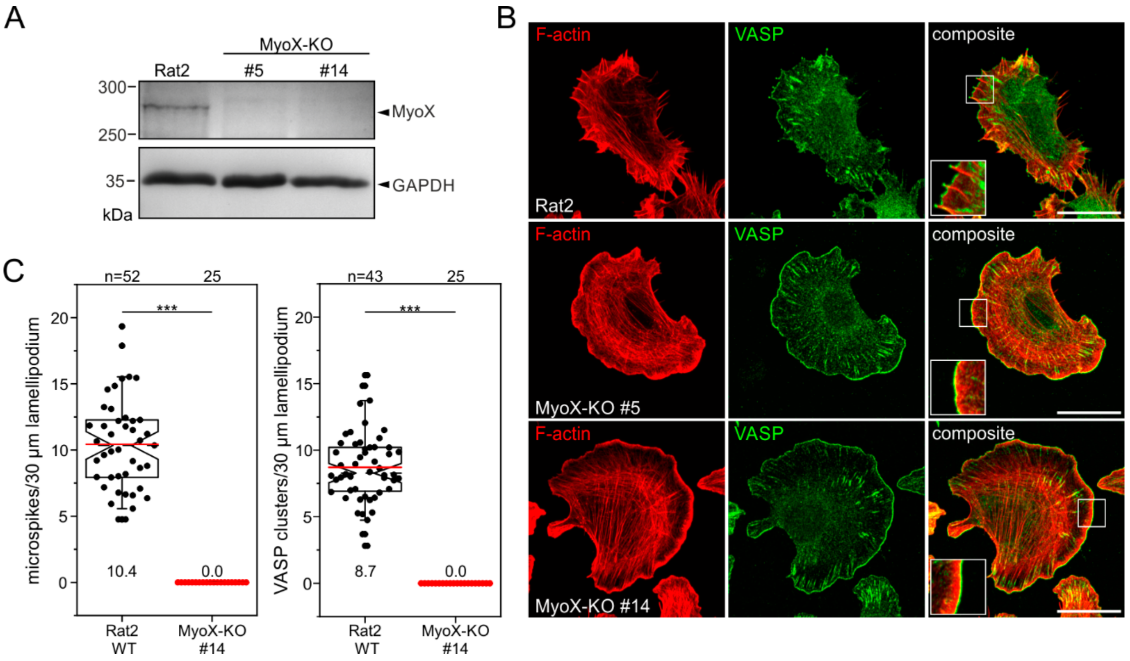


**Fig. S20. Elimination of MyoX in Rat2 cells also abolishes microspikes and VASP clustering.** (*A*) Generation of Rat2-derived mutant cell lines lacking MyoX protein expression. Immunoblot confirming the elimination of MyoX in two independent single-knockout Rat2 mutants (#5 and #14). Loading control: GAPDH. (*B*) Rat2 wild-type and the two independent MyoX-KO mutant cells stained for endogenous VASP and F-actin showing complete loss of microspikes and VASP clusters in the mutant despite prominent localization of VASP at the leading edge. Bars, 20 µm. Insets, enlarged images of boxed regions. (*C*) Quantification of microspikes and VASP clusters in wild-type and mutant cells. Notched boxes in box plots indicate 50% (25-75%) and whiskers 90% (5-95%) of all measurements; arithmetic means are highlighted in red and dashed black lines depict the medians. Statistical analysis by Mann-Whitney-U test. ***p≤0.001. n, number of analyzed cells. C shows pooled data from three independent experiments.

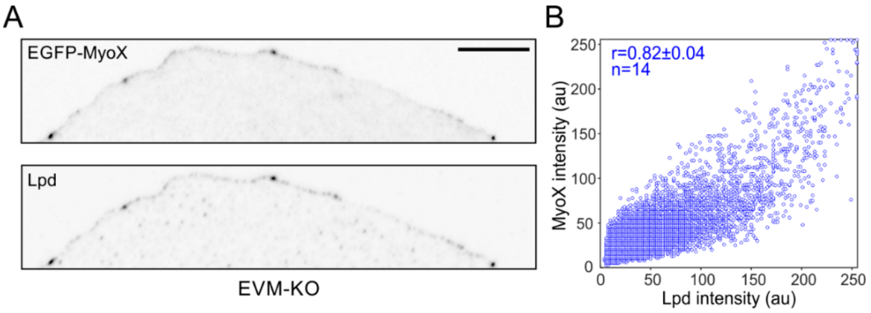


**Fig. S21. MyoX and Lpd colocalize in leading edge clusters in the absence of Ena/VASP proteins.** (*A*) Inverted confocal images of representative leading edges in Ena/VASP-deficient EVM-KO cells expressing EGFP-MyoX and additionally labelled for Lpd. Bar, 10 µm. (*B*) Cytofluorogram with respective intensity distributions and calculated Pearson correlation coefficient (r). Mean ± SD is shown; n indicates number of analyzed cells.

**Movie Legends**

**Movie S1: Loss of all I-BAR-family members perturbs lamellipodium formation in B16-F1 cells.** B16-F1 and 4xKO I-BAR cells were seeded onto laminin-coated, glass-bottom dishes and recorded by time-lapse phase-contrast imaging using a 100x objective. Note the chaotic behavior of the protruding cell front in the 4xKO I-BAR mutant. Time, min:sec. Bar, 10 µm.

**Movie S2: Combined loss of the two most abundant I-BAR-family members IRSp53 and IRTKS does not impair filopodium formation in NIH 3T3 fibroblasts.** NIH 3T3 wild-type and 2xKO I-BAR cells were seeded onto low fibronectin ($1 \mu g \cdot ml^{-1}$) on glass-bottom dishes and treated with high concentrations of the Arp2/3 inhibitor CK666 (200 µM) for 2 h, followed by time-lapse phase-contrast imaging using a 100x objective. Time, min:sec. Bar, 20 µm.

**Movie S3: Surface-bound Lpd-WT and Lpd-44A recruit and cluster VASP to drive processive actin filament elongation in the presence of capping protein.** Polymerization of 1 µM actin (10 % Atto488-labeled) on Lpd-derivatized, SNAP-Capture magnetic beads in the presence of 50 nM CP after pre-incubation with buffer or soluble VASP. Note weak actin assembly on Lpd-WT-coated beads in the absence VASP. TIRF images were captured in 3 sec intervals for a period of 15 min. Time is indicated in min:sec. Bar, 10 µm.

**Movie S4: Loss of Lpd alone or in combination with all expressed I-BAR proteins diminishes the formation of microspikes.** The cells indicated were transfected with

EGFP-LifeAct to monitor actin dynamics during migration on laminin-coated, glass-bottom dishes by time-lapse imaging using a 100x objective. Note the smooth lamellipodium after combined loss of Lpd and all expressed I-BAR family members in the 5xKO. Time is indicated in min:sec. Bar, 10 µm.

**Movie S5: The proline-rich region in the C-terminus of Abi1$^{328\text{-}481}$ is required for recruitment and clustering of VASP to allow processive actin filament elongation in the presence of CP.** Polymerization of 1 µM actin (10% Atto488-labeled) on Abi1-derivatized SNAP-Capture magnetic beads in the presence of 50 nM CP after pre-incubation with soluble VASP. In contrast to SNAP-tagged, wild-type Abi1$^{328\text{-}481}$, Abi1-mut devoid of the proline-rich sequences, was unable to recruit VASP to initiate processive actin assembly in the presence of CP. TIRF images were captured in 3 sec-intervals for a period of 15 min. Time is indicated in min:sec. Bar, 10 µm.

**Movie S6: Loss of MyoX eliminates microspikes.** B16-F1 and MyoX-KO cells were transfected with EGFP-LifeAct to monitor actin dynamics during migration on laminin-coated, glass-bottom dishes by time-lapse imaging using a 100x objective. Time is indicated in min:sec. Bar, 10 µm.

**Movie S7: Elimination of MyoX abrogates microspikes.** B16-F1 and MyoX-KO mutant cells were transfected with EGFP-fascin to monitor microspikes in cells migrating on laminin-coated, glass-bottom dishes by time-lapse imaging using a 100x objective. Time is indicated in min:sec. Bar, 10 µm.

**Movie S8: Loss of Ena/VASP proteins dramatically increases lateral movement of MyoX clusters at the protruding cell edge.** MyoX-KO and EVM-KO mutant cells were transfected with EGFP-MyoX, to monitor the dynamics of MyoX clusters at the leading edge of cells migrating on laminin-coated, glass-bottom dishes by time-lapse, TIRF imaging using a 100x objective. Note that the speed of the laterally moving MyoX clusters in reconstituted MyoX cells was still about four times slower as compared to the extremely rapidly moving MyoX clusters in EVM-KO cells. Time is indicated in min:sec. Bar, 10 μm.

**Movie S9: Loss of Ena/VASP decouples MyoX clusters from fascin-bundled microspikes.** MyoX-KO and EVK-KO mutant cells were transfected with mCherry-MyoX and EGFP-fascin to monitor accumulation of MyoX and fascin at the protruding cell front. Cells migrating on laminin-coated, glass-bottom dishes were recorded by time-lapse, dual-color TIRF imaging using a 100x objective. Bar, 5 μm. Note the lack of fascin accumulation at the base of MyoX clusters in EVM-KO cells, as opposed to reconstituted MyoX-KO cells.
